# Beckwith-Wiedemann syndrome multiomic analysis of hepatoblastoma uncovers unique tumor heterogeneity and cellular landscapes, including transition cells leading to tumor formation

**DOI:** 10.1101/2025.09.17.676802

**Authors:** Snehal Nirgude, Elisia Tichy, Yuanchao Zhang, Rose Pradieu, Michael Xie, Kathrin M. Bernt, Suzanne P. MacFarland, Jennifer M. Kalish

## Abstract

**Background:** Beckwith-Wiedemann syndrome (BWS) is an overgrowth and cancer predisposition syndrome caused by epigenetic alterations on chromosome 11p15 that predisposes children to multiple cancer types, including hepatoblastoma. Hepatoblastoma is heterogenous in nature, and the 11p15 changes that cause BWS can also be found as a somatic alteration in nonBWS hepatoblastomas, further adding complexity to this disease.

**Methods:** To understand the impact of the predisposition molecular cues in BWS hepatoblastoma, we interrogated BWS and nonBWS hepatoblastomas, as well as adjacent normal liver, using a multiomic approach [single nuclei RNA-sequencing (snRNA-seq) + single nuclei assay for transposable-accessible chromatin sequencing (snATAC-seq)].

**Results:** Our approach identified an enrichment of the WNT signaling pathway in BWS hepatoblastoma. Despite similar histology, we found greater tumor heterogeneity and embryonic transcriptional signatures in BWS hepatoblastoma. Furthermore, using pseudotime analysis, we identified a population of transition cells in BWS, with unique molecular profiles, which likely promote the precancer to cancer neoplastic transition in BWS.

**Conclusions:** This study highlights key signaling pathways, particularly WNT, and identifies a unique population of transition cells that may drive neoplastic transformation in BWS hepatoblastoma. These findings provide new insights into the molecular events leading to cancer in BWS and suggest potential targets for early intervention and prevention strategies.

## Background

Hepatoblastoma accounts for 60% of pediatric hepatic malignancies, with an incidence rate of 1.7 cases per million per year (1). This rate has increased steadily over the past several decades. Hepatoblastoma is typically diagnosed within the first five years of life (2) and is associated with multiple cancer predisposition syndromes, including Beckwith-Wiedemann syndrome (BWS) (3, 4). In patients with BWS, hepatoblastoma tends to be diagnosed earlier, often before 30 months of age (5). BWS is an overgrowth syndrome caused by epigenetic and structural alterations on chromosome 11p15 (6). Allele-specific differential methylation of two imprinting centers (IC1 and IC2) within the 11p15 region regulates the expression of a cluster of growth-related genes, including *Cyclin dependent kinase inhibitor 1C* (*CDKN1C*) and *Insulin-like growth factor 2* (*IGF2*). Specific BWS subtypes are defined by the methylation status of these imprinting centers. However, BWS patients with IC2 loss of methylation (IC2 LOM, reduced *CDKN1C* expression) or paternal uniparental isodisomy of chromosome 11 (pUPD11, reduced *CDKN1C* expression and increased *IGF2* expression) are more likely to develop hepatoblastoma (6).

Hepatoblastoma is a histologically and molecularly heterogeneous tumor, broadly classified into two main categories: epithelial and mixed epithelial–mesenchymal types (7). More than 50% of hepatoblastomas are of epithelial origin (8) and encompass several histologic subtypes (7). Among these, the fetal subtype is associated with a more favorable overall prognosis, while the small cell undifferentiated subtype is linked to significantly poorer outcomes (9). Mixed-type hepatoblastomas show a mesenchymal pattern and teratoid features, suggesting a multidirectional trajectory of hepatoblastoma development (10). These histologic subtypes have distinct clinical implications—for instance, children with pure fetal histology can achieve long-term survival with surgical resection alone (11) (12). In contrast, patients with more aggressive subtypes, such as mixed epithelial–mesenchymal, require multiple rounds of chemotherapy with significant short and long term toxicity, as illustrated by a reported case of BWS-associated hepatoblastoma (13). These observations highlight the importance of studying the hepatoblastoma-associated tumor heterogeneity in predicting clinical outcomes. Many groups have defined hepatoblastoma tumor heterogeneity transcriptomically, using their own criteria (14–23); however, BWS-driven hepatoblastoma tumor heterogeneity has never been examined at single-nucleus resolution.

At the genomic level, hepatoblastoma is driven by a constellation of molecular alterations, including CTNNB1 (β-catenin) mutations, 11p15 abnormalities, and other oncogenic drivers (18). However, the chronological order and functional interdependence of these events remain poorly defined. Pilet *et al.* (24), provided evidence that 11p15 alterations precede CTNNB1 mutations in sporadic (nonBWS) hepatoblastoma, suggesting that epigenetic disruption at 11p15 may serve as an early oncogenic event that establishes a tumor-permissive molecular landscape. We previously addressed this hypothesis, in part, by defining predisposition and oncogenic gene expression signatures in BWS hepatoblastoma using bulk transcriptomic analysis (3). That work revealed that BWS tumor-adjacent liver tissue, though histologically normal, carries gene expression profiles indicative of early neoplastic transition, implicating the presence of a latent predisposition state. In a subsequent study, we used single-nucleus RNA sequencing to profile tumor-adjacent liver tissues from both BWS and nonBWS patients, identifying a PPARA-driven metabolic signature uniquely enriched in BWS livers (4). These findings led us to propose a model wherein 11p15 epigenetic alterations induce metabolic reprogramming, creating a precancerous molecular environment that contributes to hepatoblastoma predisposition in BWS (4). However, that study did not characterize the full cellular and molecular trajectory from predisposed BWS liver to overt hepatoblastoma.

The development of tumors from normal tissue typically proceeds through a multistep process, where the precancerous or transition state serves as a key intermediate (25). These transition-state cells exhibit hybrid transcriptional identities, reflecting partial features of both normal and malignant states (26). While such cells play essential roles in tissue remodeling during development, they are also implicated in drug resistance, cancer initiation, and metastasis (27). Pseudotime analysis of single-cell data has been widely used to infer these intermediate states along developmental or oncogenic trajectories (25, 26, 28, 29).

Given the distinct oncogenic underpinnings of BWS hepatoblastoma (3), the current study employed a multiomic single-nucleus approach to investigate the cellular architecture, molecular pathways, tumor heterogeneity, and developmental trajectory of BWS-associated hepatoblastoma. We compared these features with those of nonBWS hepatoblastoma to delineate shared and unique tumorigenic mechanisms. Using pseudotime trajectory inference, we further identified transition cell populations that appear to mediate the shift from precancerous to malignant states and demonstrated that these transition cells are molecularly distinct in BWS, relative to nonBWS cases.

## Methods

### Patients and samples

BWS1, BWS2, BWS3, and BWS4 represent the BWS liver/tumor-adjacent cohort; BWST1, BWST2, BWST3 and BWST4 represent the BWS hepatoblastoma cohort; NS1, NS2, NS3 represent the nonBWS liver/tumor-adjacent cohort having no cancer predisposition syndrome; and NST1, NST2, NST3 represent the nonBWS hepatoblastoma cohort. The methylation profile for both the BWS and nonBWS hepatoblastoma and tumor-adjacent liver (4) were confirmed by pyrosequencing (**Supplementary Fig. 1, Supplementary Table 1**). Clinical genetic testing for BWS molecular characterization was performed in blood at the University of Pennsylvania Genetic Diagnostic Laboratory, as previously described (30). All tumor-adjacent liver samples were collected from patients during their surgical hepatoblastoma resection and were confirmed as normal adjacent tissue beyond the tumor margins by a clinical pathologist. All the samples were snap-frozen in liquid nitrogen and stored at −80°C.

### Histology of livers

nonBWS tumor-adjacent liver/NS, nonBWST/NST (hepatoblastoma), BWS tumor-adjacent liver and BWST (hepatoblastoma) samples were FFPE processed, sectioned, and stained using standard hematoxylin and eosin staining protocols. Sections were imaged using Leica Aperio AT2 slide scanner.

### Pyrosequencing

Pyrosequencing was performed on bisulfite-converted DNA using the Pyromark PCR kit (Qiagen), and sequencing was performed on a Pyromark Q48 Autoprep (Qiagen), as described (4). Sequencing primers are listed in the study (**Supplementary Table 2**) (4).

### Single nuclei isolation, library preparation and sequencing

The nuclei from liver/tumor-adjacent and tumor samples belonging to both BWS and nonBWS cohorts were isolated as described (4). 10X Chromium libraries were prepared using the 10x Genomics Chromium Single Cell Multiome Assay for Transposase-Accessible Chromatin (ATAC) and Gene Expression Reagent kit v1, according to the manufacturer’s protocol by CHOP’s Center for Applied Genomics (CAG) core. A single sequencing run was performed to avoid batch effect. Gene expression libraries were uniquely indexed using the Chromium dual Index Kit, pooled, and sequenced on an Illumina NovaSeq 6000 sequencer in a paired-end, dual indexing run, targeting 20,000 mean reads per cell. Data was then processed using the Cell Ranger pipeline (10x Genomics, version 3.1.0) for demultiplexing and alignment of sequencing reads to the (reference hg38) transcriptome, as described (4).

### Single nuclei-RNA sequencing

#### 1. snRNAseq data preprocessing, quality control and data integration

The data was processed using Seurat package (version 4.9) (31–34) in R (version 4.3), as described (4). The nuclei with transcript counts outside the 200-50000 range, >5% mitochondrial reads in snRNA-seq data, were excluded (4). The DoubletFinder R package (version 2.0.3) was used for identifying and removing doublets (4) (35).

After performing initial quality control (QC) as described (4), snRNAseq data was consolidated into a unified count matrix using basic R Matrix.utils. The data was integrated using Seurat’s built-in functionalities. The dataset was subject to normalization with “NormalizeData” followed by data scaling (”ScaleData” function), as described (4). “FindVariableFeatures” was used to select the top 2000 features using “vst” method. Principal Component Analysis (PCA) was conducted with “RunPCA” function and top 30 principal components were retained for dimensional reduction (4). Batch effects were removed using Harmony (36) with all default parameters and maximum 30 iterations using the “RunHarmony” function, followed by integration (4).

#### 2. snRNAseq clustering, marker genes and cell type identification

The harmonized data was subject to cell clustering via Seurat’s original Louvain clustering algorithm (4). The data was subject to “RunUMAP”, followed by “FindNeighbors” and finally “FindClusters” at a resolution level of 0.5 was employed, as described (4). The top 50 marker genes for each cluster were then studied (4).

For our tumor heterogeneity study and pathway study, we used the gene list in **Supplementary Data File 1, 2** and subjected it to the AddModuleScore feature from Seurat package. The score was calculated to study the distribution of the expression of the genes associated with a given category of tumor classification and the score was plotted as box plot. A nonparametric statistics test “Wilcoxon Rank Sum test” was used to rank gene expression significance due to the unique nature of the single cell RNA-seq. Any gene deemed as “Marker gene” would have minimal expression fold change at 0.1 (log-scale).

Cell type identification was carried out primarily using marker genes, as described (4). We aggregated the list cluster marker genes from a literature search and then we utilized two R packages SingleR (version 1.0.1) (37) and ScType (available at https://github.com/IanevskiAleksandr/sc-type) (38) (4).

#### 3. snRNAseq pathway analysis comparing tumor clusters between BWS and nonBWS cohorts

For pathway analysis, we used the Single Cell Pathway Analysis (SCPA) R package v1.6.2 (39) to characterize the gene functional enrichment analysis within a given combination of condition (BWS/BWST/NS/NST) and cluster, as described (4). The gene functional annotation was collected from the Molecular Signatures Database (MSigDB) (40, 41) by the Broad Institute, which includes curated gene sets (4). We used the R package msigdbr v7.5.1 to get the MSigDB gene sets. Our analysis primarily focused on the curated gene set collections: “C2 – the curated gene sets”. SCPA uses a multivariate distribution testing approach to identify pathways enriched in a given combination of condition and cell type and provides transcriptional changes for a given pathway within each combination of condition and cluster (4). We employed the Benjamini-Hochberg (BH) correction procedure to reduce multiple testing errors.

### Single-nuclei ATACseq data analysis

For quantification of single nuclei ATACseq data, we first constructed a cell-by-peak matrix using PICsnATAC (https://github.com/Zhen-Miao/PICsnATAC). Then we used Seurat R package (v5.0) and harmony package (version 1.2.3) to perform dimension reduction and data integration. Cell type labels obtained from snRNAseq were transferred to ATACseq based on cell barcodes. Cells within each cell type were aggregated to generate bulk ATACseq-type data using the “addGroupCoverages” function in ArchR (v1.03) (4). Chromatin accessibility peaks were identified using the “addReproduciblePeakSet” function, utilizing the MACS2 peak identification pipeline with the human genome build hg38 as a reference. Within each cell type, we directly compared syndromic cells (BWS) to nonsyndromic cells (NS) using archR function “getMarkerFeatures”. We collected all peaks with p-value less than 0.05 and logFC greater than 1 (4). Peaks within each cell groups were counted and annotated using data from EnsDb.Hspiens.v86 (R package version 2.99.0) (4). The resulting figure was plotted using ggplot2 (4).

### Virtual copy-number profiles

The virtual copy-number profiles were predicted using the raw snRNA-seq counts of the 147,820 nuclei to distinguish tumor from normal nuclei. For this purpose, we used the R package InferCNV v1.20.0 (42) with the workflow and parameters suggested by the inferCNV tutorial at https://github.com/broadinstitute/infercnv/wiki/Running-InferCNV, and we used healthy hepatocytes (Cluster 0) and cholangiocytes (Cluster 6) as reference nuclei. In the results of InferCNV, the CNV states that were predicted by the InferCNV i6 Hidden Markov Model (HMM) and filtered by the InferCNV Bayesian mixture model were used for visualizations. The InferCNV i6 HMM has the following states: state 1 represents complete loss; state 2 represents loss of one copy; state 3 represents neutral; state 4 represents addition of one copy; state 5 represents addition of two copies; state 6 represents addition of three or more copies but modeled as addition of three copies. The CNV states of all genes and cells were visualized using a heatmap. The average CNV states of the genes on the chromosome 11 of the cells in each sample were also visualized using a heatmap. The number of genes with non-neutral CNV states in each cell were visualized using boxplots.

### Pseudotime analysis

The R package slingshot v2.12.0 (43) was used for pseudotime analysis to infer the trajectory of BWS hepatoblastoma. The pseudotime and trajectory were inferred from the PCA embeddings of a subset of clusters, including hepatocyte cluster and tumor clusters 1-5, with the hepatocyte cluster as the starting cluster. To identify the transition cell population, we used the pseudotime binning approach as described (44). Based on the changes of the number of genes with non-neutral CNV states in each cell across 6 equal pseudotime bins, we defined three cell states: precancer (pseudotime range from 0 to 24), transition (pseudotime range from 24 to 34) and cancer (pseudotime range from 34 to the maximum). We then performed marker gene analysis as described above by comparing transition vs precancer states and cancer vs transition states. Further we analyzed the pathways enriched in transition and cancer states by applying Gene Set Enrichment Analysis (GSEA), implemented in the R package fgsea 1.30.0 (45) on the log2 fold changes of transition vs precancer states and cancer vs transition states.

### Integrated snRNA-seq and snATAC-seq analysis using SCENIC plus

The snRNA-seq and snATAC seq data was integrated using single-cell multiomic inference of enhancers and gene regulatory networks (SCENIC +) (46), as described (4). The python package SCENIC+ v1.0a2 was used to infer both transcription factor activity and candidate enhancer regions and regulated genes network (eRegulons) (4). We used the combinations of conditions and pseudotime-defined precancer, transition and cancer states to group the hepatocyte cluster and the tumor clusters 1-5 for the SCENIC+ analysis (4). We used the python package pycisTopic v2.0a0 (46) to process snATAC-seq data and identify sets of co-accessible regions (termed topics) and differentially accessible regions (DARs) (4). We created a cisTarget database using the pycisTopic results and the python package create_cisTarget_databases versioned at the Git commit 3b62de8 (37) (4). We used the python package pycistarget v1.1 (46) for motif enrichment analysis. Using the snRNA-seq data, DARs, topics, and the enriched motifs, eRegulons were inferred by SCENIC+. From the SCENIC+ inferred eRegulons, high-quality eRegulons were selected for visualizations, by adapting the eRegulon filtering procedure in the SCENIC+ tutorial at https://github.com/aertslab/scenicplus/blob/v1.0.0/notebooks/pbmc_multiome_tutorial.ipynb (37) (4). For the selected high-quality eRegulons, the log(1+x) transformed average normalized counts of the transcription factors and the SCENIC+ Regulon Specificty Scores (RSS) were visualized using a heatmap (4). From the selected high-quality eRegulons, another subset of eRegulons was selected based on their transcription factor expression levels in the transition cells, which includes CTNNB1_direct_+/+, ETV1_direct_+/+, LEF1_direct_+/+, PLAG1_direct_+/+, RFX3_direct_+/+, ZNF518A_direct_+/+, and ZNF704_direct_+/+. For the eRegulons selected based on their transcription factor expression levels in the transition cells, an interaction network was created to only include the eRegulon gene targets that are involced in Kegg_ECM_receptor_interaction pathway and Kegg_Gap_Junction pathway, by adapting the eRegulon network creation procedure in the SCENIC+ tutorial. The interactions in the interaction network between the selected eRegulon transcription factors, target genes, and target regions are visualized using a SCENIC+ network plot.

### Ligand-receptor interactions

The ligand-receptor interaction analysis was performed using the R package CellChat v2.1.2 (47) with the workflow and parameters suggestedby the CellChat tutorial at https://github.com/jinworks/CellChat/blob/v2.1.2/tutorial/CellChat-vignette.Rmd. The human CellChatDB was used for the analysis. The pseudotime-defined cell states were used as cell groups to analyze ligand-receptor interactions. The p-values and communication probabilities of the ligand-receptor interactions were visualized using a dot plot.

### Results Clinical cohort

One aim of this study was to decipher the cell-type-specific molecular signatures driving BWS hepatoblastoma and to determine whether these signatures were distinct from those of nonBWS hepatoblastoma samples. To address this, we used matched tumor samples from the same set of patients as described in our previous BWS tumor-adjacent liver study (4). The samples were derived from 4 BWS patients and 3 nonBWS patients, as described (4). NonBWS samples were age and sex stratified with our BWS samples, based on sample availability within the institutional biobank. Histologic review of both tumor-adjacent tissue and tumor tissue revealed similar liver and tumor gross morphologies within BWS and nonBWS groups (**Supplementary Fig. 2 and Supplementary Fig. 3**). **Supplementary Table 3** displays the clinical information for both cohorts, including the histological diagnoses assigned by the pathologist. We also confirmed the methylation status of the two imprinted regions on chromosome 11p15 (*H19*/*IGF2*:IG-DMR and *KCNQ1OT1*:TSS-DMR) in both nonBWS and BWS tumor samples by performing pyrosequencing for *KCNQ1OT1*:TSS-DMR (IC2) and *H19*/*IGF2*:IG-DMR (IC1) (**Supplementary Fig. 1a-d**). The nonBWS tumor samples exhibited normal methylation of IC2; however, one nonBWS hepatoblastoma tumor sample did exhibit a gain of methylation at IC1. All BWS tumor samples displayed aberrant methylation at both IC1 and IC2, as were previously reported (3). **Supplementary Table 1** shows methylation profile for nontumor/tumor-adjacent tissue, adapted from (4).

### The cellular landscape of BWS livers, BWS tumors, nonBWS liver and nonBWS tumors

We conducted an integrated multiomic study (single-nucleus RNA and ATAC sequencing) of hepatoblastoma and tumor-adjacent tissues from both BWS and nonBWS patients. Single-nucleus RNA sequencing (snRNA-seq) of BWS (tumor-adjacent and tumor) and nonBWS (tumor-adjacent and tumor) samples revealed 18 transcriptionally distinct clusters, encompassing all expected normal liver cell types and several tumor-specific clusters (**Fig. 1a, Supplementary Table 4**), based on our previously established snRNA-seq analysis pipeline (4). Following quality control, including the removal of low-quality nuclei and putative doublets, 147,820 nuclei were retained for downstream analysis. Clustering showed minimal inter-sample batch variation (**Fig. 1b**). Cell type annotation was performed using a combination of SingleR, ScType, and manual annotation methods. Marker genes were identified using Seurat’s *FindMarkers* function, as described (4) (**Supplementary Data File 3** – **Top 50 cell markers in each cluster**). Liver cell types—including hepatocytes, cholangiocytes, endothelial cells, hepatic stellate cells, immune cells, and Kupffer cells—were annotated using previously validated markers (4) (**Fig. 1c**). Five tumor-specific clusters (tumor clusters 1–5) were characterized by high expression of known hepatoblastoma markers such as *GPC3* (48, 49), *MEG3* (18, 21, 50) and *MEG8* (18, 50) (**Fig. 1c**). All 18 clusters were represented across every patient sample (**Fig. 1d**), with tumor clusters being predominantly derived from tumor tissues (**Fig. 1e**). To complement the transcriptional data, we performed snATAC-seq and annotated chromatin accessibility profiles using Paired-Insertion Counting snATAC (PICsnATAC) (51), Seurat, and label transfer from our snRNA-seq dataset. We identified 18 chromatin-accessibility-based cell populations across all samples from both cohorts (**Supplementary Fig. 4a**). Tumor versus tumor-adjacent comparisons revealed distinct chromatin accessibility patterns between BWS and nonBWS tumors (**Supplementary Fig. 4b**), suggesting divergent gene regulatory landscapes in BWS-associated versus sporadic hepatoblastoma. Enrichment of snATAC-seq marker peaks across all annotated cell types and conditions (BWS and nonBWS, tumor and tumor-adjacent) was quantified and visualized (**Supplementary Fig. 4c and 4d**). These peaks mapped to UTR, promoter, distal, intronic, and exonic regions, consistent with expected regulatory element distributions.

**Figure 1:**
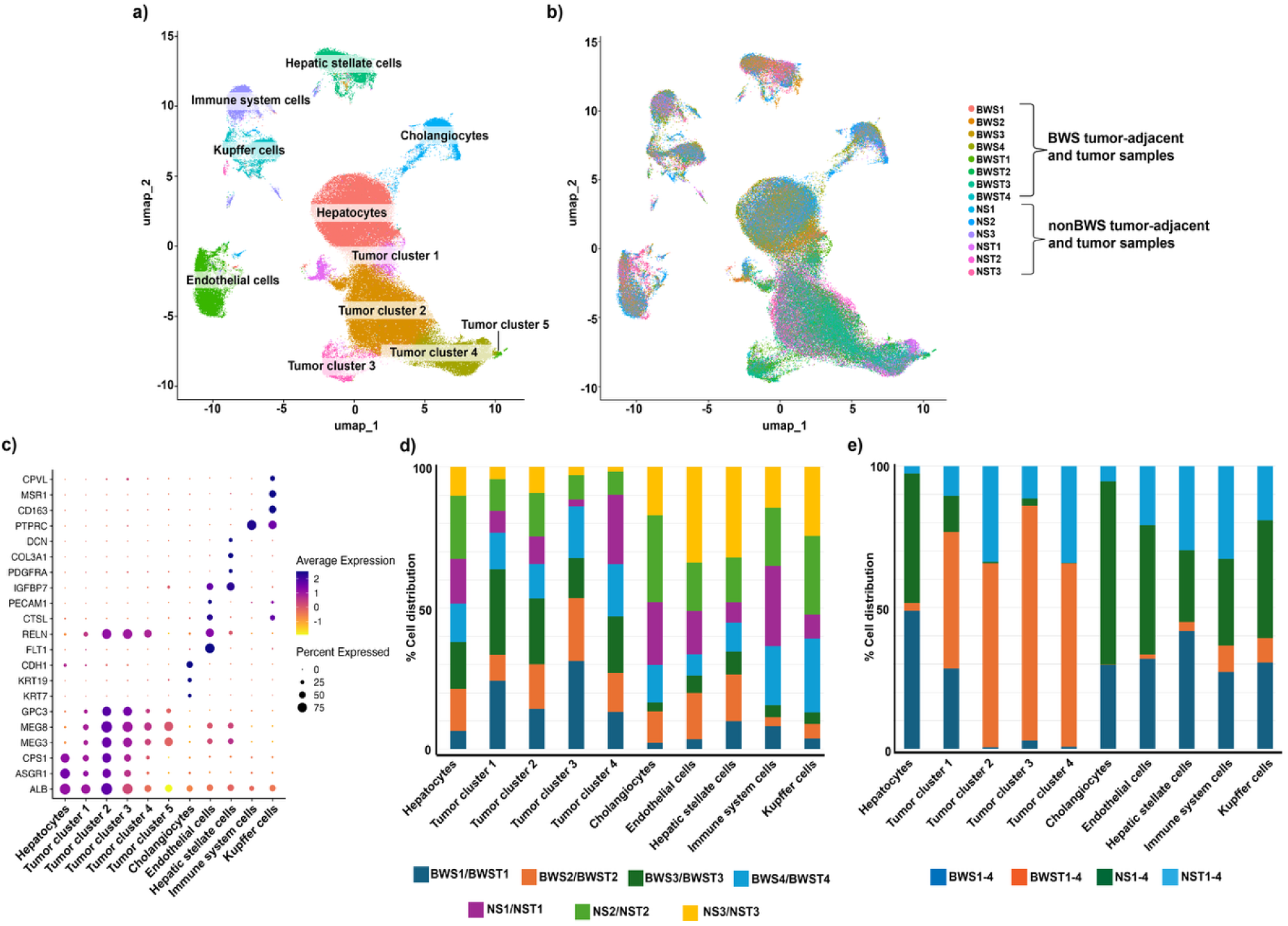
snRNA-seq profile of liver/tumor-adjacent and hepatoblastoma samples belonging to BWS and nonBWS cohorts. (**a**) Uniform manifold approximation and projection (UMAP) visualization of 147,820 nuclei isolated from 7 liver and hepatoblastoma samples. This single-nuclei transcriptome atlas identified major liver cell types including hepatocytes, cholangiocytes, hepatic stellate cells, Kupffer cells, endothelial cells, immune system cells and tumor cell clusters in both BWS and nonBWS sample cohorts. (**b**) UMAP of BWS-liver (BWS) (n=4), BWS-Tumor (BWST) (n=4), nonBWS-liver (NS) (n=3) and nonBWS-Tumor (NST) (n=3) using snRNA-seq demonstrates that both liver groups contain the same cell types. (**c**) Dot plots of snRNA-seq dataset showing gene expression patterns of cluster-enriched markers for liver and tumor cell types. The diameter of the dot corresponds to the proportion of cells expressing the indicated gene and the density of the plot corresponds to average expression relative to all cell types. **(d)** Stacked column graphs displaying the contribution of BWS and nonBWS patients to each cell population. The percentages from each individual color add up to the total number of cells in that sample type. **(e)** Stacked column graphs displaying the contribution of each cell population belonging to BWS and nonBWS sample types to normal and tumor. The percentages from each individual color add up to the total number of cells in that sample type.

To confirm that the transcriptionally defined tumor clusters indeed represented malignant cells, we performed inferCNV analysis, using hepatocytes and cholangiocytes as reference populations. This analysis revealed widespread copy number variations (CNVs) in tumor clusters from both BWS and nonBWS cohorts compared to reference cells (**Supplementary Fig. 5a**). consistent with their neoplastic identity. Given that BWS is associated with epigenetic and genetic alterations on chromosome 11, we further investigated CNVs across chromosome 11. Distinct profiles were observed among the four groups—BWST (BWS tumor), BWS (BWS tumor-adjacent), NST (nonBWS tumor), and NS (nonBWS tumor-adjacent)—further validating our tumor classification strategy (**Supplementary Fig. 5b**).

### Pathway analysis reveals enrichment of distinct molecular signaling in BWS hepatoblastoma

Following the definition of the cellular landscape in BWS and nonBWS hepatoblastoma, we next investigated the molecular signaling pathways enriched in tumor tissue compared to tumor-adjacent (non-tumor) tissue in both cohorts. To this end, we performed differential pathway analysis between hepatocytes and tumor clusters 1–5 within each group. Pathways enriched across all tumor clusters collectively are shown in **Fig. 2**. We observed distinct enrichment profiles between BWST and nonBWST (**Fig. 2a**, **2b**). Specifically, BWST clusters demonstrated enrichment for WNT_Signaling, Gap_Junction, and GnRH_Signaling pathways based on KEGG gene lists (**Supplementary Fig. 6, Supplementary Data File 4**). In contrast, nonBWST clusters were predominantly enriched in Oxidative_Phosphorylation, Fatty_Acid_Metabolism, and ECM_Receptor_Interaction pathways (**Supplementary Fig. 6, Supplementary Data File 4**). To further investigate these distinctions, we calculated module scores for the aforementioned pathways and compared their expression profiles between hepatocytes and tumor clusters across both BWS and nonBWS groups (**Fig. 2c-h**). WNT signaling was elevated in both cohorts, with a stronger enrichment observed in the BWST group, whereas fatty acid metabolism was more prominently upregulated in the non-BWST group. These results demonstrate that BWS and nonBWS hepatoblastomas harbor distinct pathway enrichment profiles, reinforcing the presence of unique tumor heterogeneity in each cohort.

**Figure 2:**
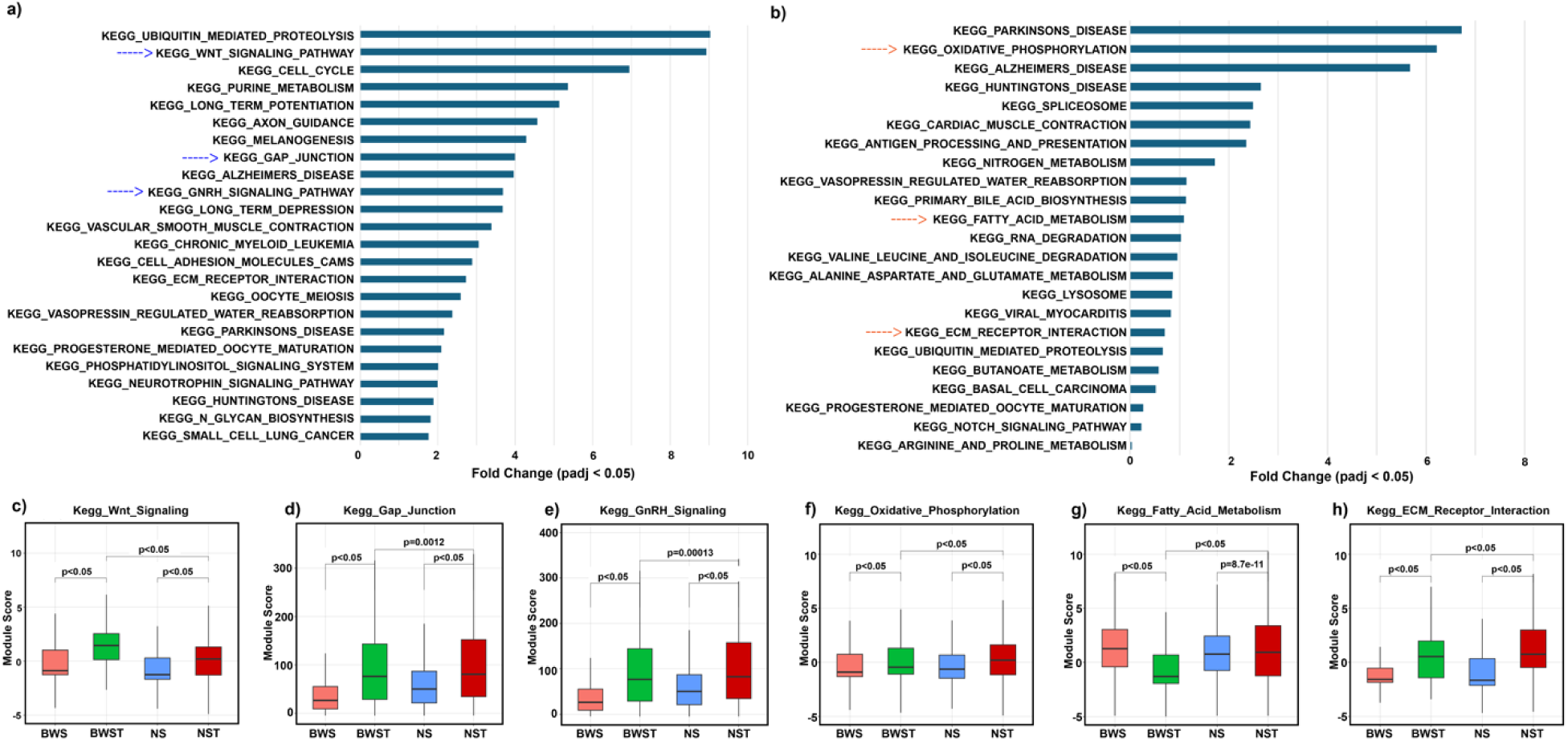
Pathway enrichment analysis of tumor clusters representing BWST and nonBWST samples. (**a**) Kegg pathways enriched in the tumor clusters of BWST when compared to hepatocyte clusters of BWS. The blue arrows indicate the pathways enriched in all the tumor clusters in BWS samples. (**b**) Kegg pathways enriched in the tumor clusters of nonBWST/NST compared to hepatocyte clusters of nonBWS/NS. The orange arrows indicate the pathways enriched in all the tumor clusters in nonBWS samples. (**c-e**) Module score for common pathways upregulated in tumor clusters of BWST when compared to hepatocyte clusters of BWS. (**f-h**) Module score for common pathways upregulated tumor clusters of nonBWST/NST compared to hepatocyte clusters of nonBWS/NS.

Having identified cohort-specific signaling features, we next sought to determine whether common molecular signatures of hepatoblastoma were preserved across both BWS and nonBWS tumors. For this, we examined the expression of the 14q32 gene signature, a well-established marker of hepatoblastoma aggressiveness (50) This signature, associated with WNT pathway activation and poor prognosis, is composed of a cluster of genes listed in **Supplementary Data File 1**. Our analysis revealed that the 14q32 signature was significantly upregulated in both BWST and nonBWST relative to their respective tumor-adjacent tissues (**Supplementary Fig. 7a**), with stronger enrichment observed in the nonBWST group. These findings highlight common molecular pathways involved across both cohorts, with differences in magnitude and context, confirming the 14q32 signature as a shared molecular hallmark of hepatoblastoma regardless of BWS status.

In addition, we queried the Reactome pathway database to further identify shared molecular features between cohorts (**Supplementary Data File 4 - Kegg and Reactome pathways enriched in cell clusters**). Notably, ROBO1 signaling emerged as another commonly enriched pathway in both BWS and nonBWS hepatoblastomas. ROBO1, a known oncogenic driver and therapeutic target in hepatocellular carcinoma (52–54). was expressed at higher levels in tumor clusters than in hepatocytes in both cohorts. This was confirmed by computing module scores for ROBO1 signaling pathway genes, which were significantly elevated in BWST and nonBWST compared to their respective non-tumor cells (**Supplementary Fig. 7b**). Interestingly, nonBWS hepatoblastoma was more highly enriched compared to BWS hepatoblastoma in ROBO1 signaling, once again demonstrating that BWS hepatoblastoma is different from other hepatoblastoma. All together, these analyses highlight that while BWS hepatoblastoma is characterized by greater enrichment of signaling pathways such as WNT and GnRH signaling, it also shares canonical hepatoblastoma features, including 14q32 signature overexpression and ROBO1 pathway activation, with nonBWS tumors, albeit at different levels. This dual profile underscores both the unique tumor biology and shared molecular underpinnings of BWS-associated hepatoblastoma.

### Transcriptomic Heterogeneity of BWS and nonBWS hepatoblastoma cells

To further dissect tumor heterogeneity and contextualize our previous pathway enrichment findings, we sought to characterize BWS and nonBWS tumor cells (BWST and nonBWST, respectively) by mapping them to established hepatoblastoma molecular classifications. To achieve this, we subsetted hepatocytes and tumor clusters (clusters 1–5) from our Seurat object and conducted pairwise comparisons between: (i) BWS hepatocytes vs. BWST, (ii) nonBWS hepatocytes vs. nonBWST, and (iii) BWST vs. nonBWST. We utilized a curated panel of hepatoblastoma gene signatures from published literature (**Supplementary Data File 2**) for this comparative analysis.

We first applied the hepatoblastoma gene expression signatures defined by Cairo *et al.* (17), which stratified tumors into two molecular subtypes: C1 (intermediate risk) and C2 (high risk/poor prognosis). In our snRNA-seq dataset, the C2 gene signature was more prominently expressed in BWST relative to adjacent non-tumor BWS hepatocytes, while both C1 and C2 signatures were comparably expressed in nonBWST (**Fig. 3a**). Furthermore, BWST displayed diminished enrichment of the C1 signature and enhanced C2 expression when compared directly to nonBWST, suggesting a more aggressive molecular profile in BWS-associated hepatoblastoma. Next, using gene sets defined by Roehrig *et al.* (18) which categorize hepatoblastoma cells as *hepatocytic*, *progenitor*, or *mesenchymal*, we observed that BWST cells exhibited increased expression of progenitor-associated genes and reduced hepatocytic signature expression compared to BWS tumor-adjacent hepatocytes. Notably, BWST also showed greater mesenchymal gene signature enrichment than BWS tumor-adjacent tissue— captured through our single-nuclei resolution approach. In contrast, nonBWST tumors expressed all three gene categories more strongly than their nonBWS tumor-adjacent counterparts (**Fig. 3b**). When comparing BWST to nonBWST, both showed similar levels of progenitor gene expression; however, nonBWST demonstrated greater enrichment of hepatocytic and mesenchymal markers, underscoring additional layers of tumor heterogeneity in BWS hepatoblastoma.

**Figure 3:**
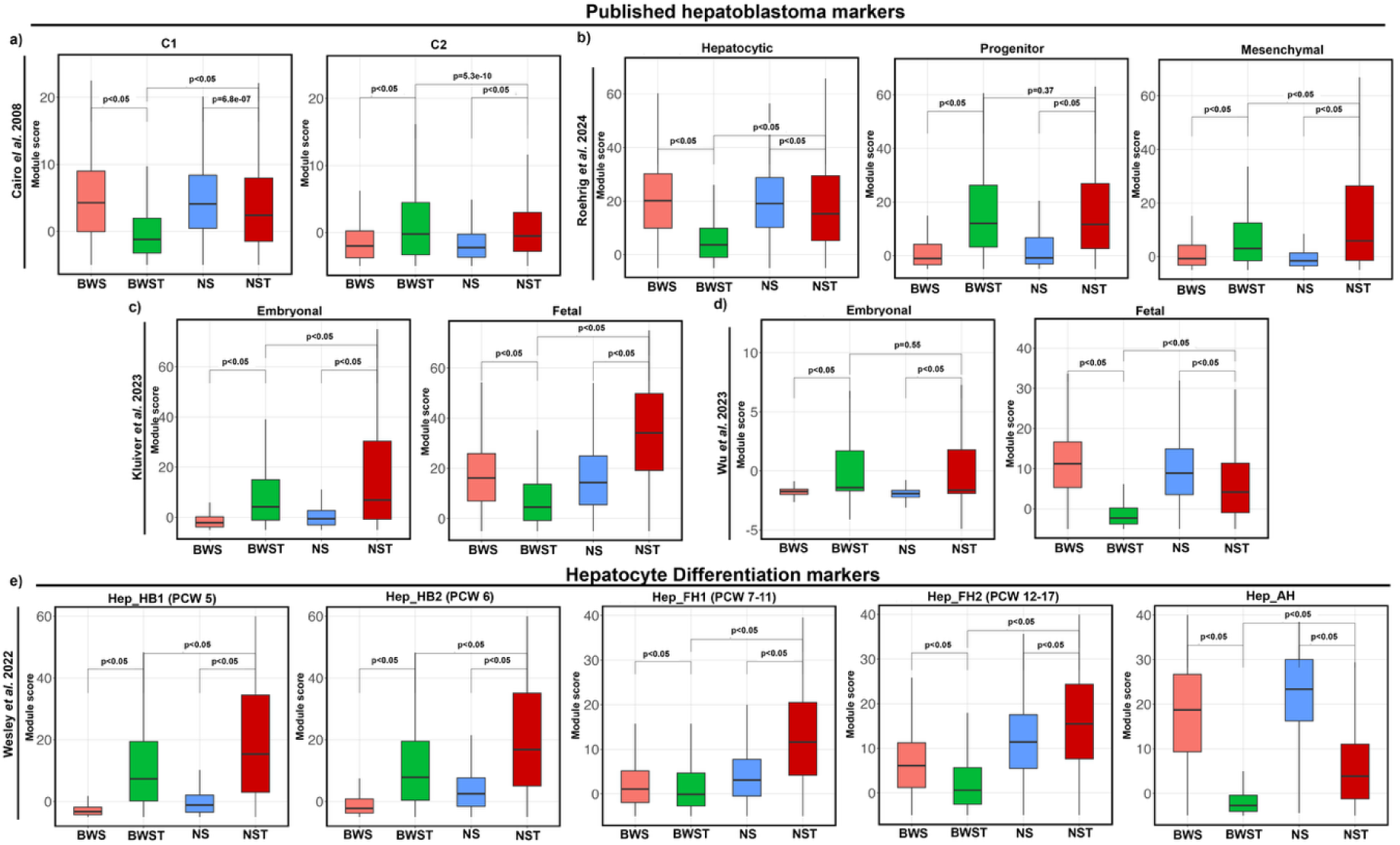
Tumor cell analysis reveals transcriptional heterogeneity in BWS and nonBWS hepatoblastoma. Box plots showing module score of published gene signatures of hepatoblastoma from single-cell RNA sequencing and bulk gene expression analyses (gene lists in Supplementary Data) in tumor-adjacent and tumor samples of BWS and nonBWS cohorts. (**a**) C1, C2 gene signatures by Cairo *et al*., (**b**) Hepatocytic, Progenitor, Mesenchymal gene signatures from Roehrig *et al*., (**c**) Embryonal and fetal gene signatures from Kluvier *et al*., (**d**) Embryonal and fetal gene signatures from Wu *et a*l., (**e**) Hepatocyte differentiation markers from Wesley *et al*. (PCW-Post conception week, Hep_HB1-Hepatoblast 1, Hep_HB2 – Hepatoblast 2, Hep_FH1 – Fetal Hepatocyte 1, Hep_FH2 - Fetal Hepatocyte 2, Hep_AH – Adult Hepatocyte).

We further examined the ‘embryonal’ and ‘fetal’ gene signatures established by Kluvier *et al.* (14) and Wu *et al.* (15). Fetal hepatoblastomas are associated with lower proliferative indices and favorable surgical outcomes, while embryonal types are more aggressive and histologically complex, often displaying epithelial-to-mesenchymal transition (EMT) and elevated AFP (14, 15). BWST cells were enriched for embryonal gene signatures compared to BWS tumor-adjacent cells. NonBWST cells; however, expressed both fetal and embryonal signatures robustly (**Fig. 3c**, **3d**). Direct comparison of the two tumor groups revealed reduced fetal marker expression in BWST relative to nonBWST, while embryonal markers were more selectively enriched in BWST. These findings suggest that BWS-associated hepatoblastomas skew more strongly toward an embryonal phenotype, whereas nonBWS tumors express a mixture of both developmental states.

Given that hepatoblastoma originates from hepatic precursors and recapitulates developmental programs (7), we assessed the developmental maturity of tumor cells by referencing single-cell RNA-seq data from normal fetal liver at varying post-conception weeks (PCWs) (55). BWST cells exhibited increased expression of hepatoblast markers associated with early developmental stages (PCW 5–6), relative to BWS tumor-adjacent tissue (**Fig. 3e**). A similar trend was seen in nonBWST samples. However, at later developmental stages (PCW 7–11 and 12–17), BWST cells lacked significant enrichment, in contrast to nonBWST, which displayed clear enrichment of fetal hepatocyte signatures from these later stages. Neither BWST nor nonBWST expressed adult hepatocyte signatures, ruling out normal liver contamination. These data suggest that BWST cells may represent an earlier developmental state than nonBWST, supporting the concept of increased cellular immaturity and altered tumor evolution in BWS hepatoblastoma.

To systematically define the molecular characteristics of BWS tumor heterogeneity, we defined two distinct gene expression modules—BWSTI and BWSTII, derived from selected hepatoblastoma signatures (**Supplementary Data File 2**). BWSTI is associated with liver progenitor features and cell proliferation, including genes such as *TOP2A, POLQ,* and *MKI67* (20). BWSTII is marked by embryonal and mesenchymal programs, with strong expression of *VIM, NKD1,* and *TNFRSF19* (14). (**Fig. 4a**, **4b**). Together, these results show that BWST cells exhibit a molecular identity characterized by early developmental origin, high proliferative potential, and mesenchymal/embryonal traits—underscoring a distinct tumor biology compared to nonBWS hepatoblastoma.

**Figure 4:**
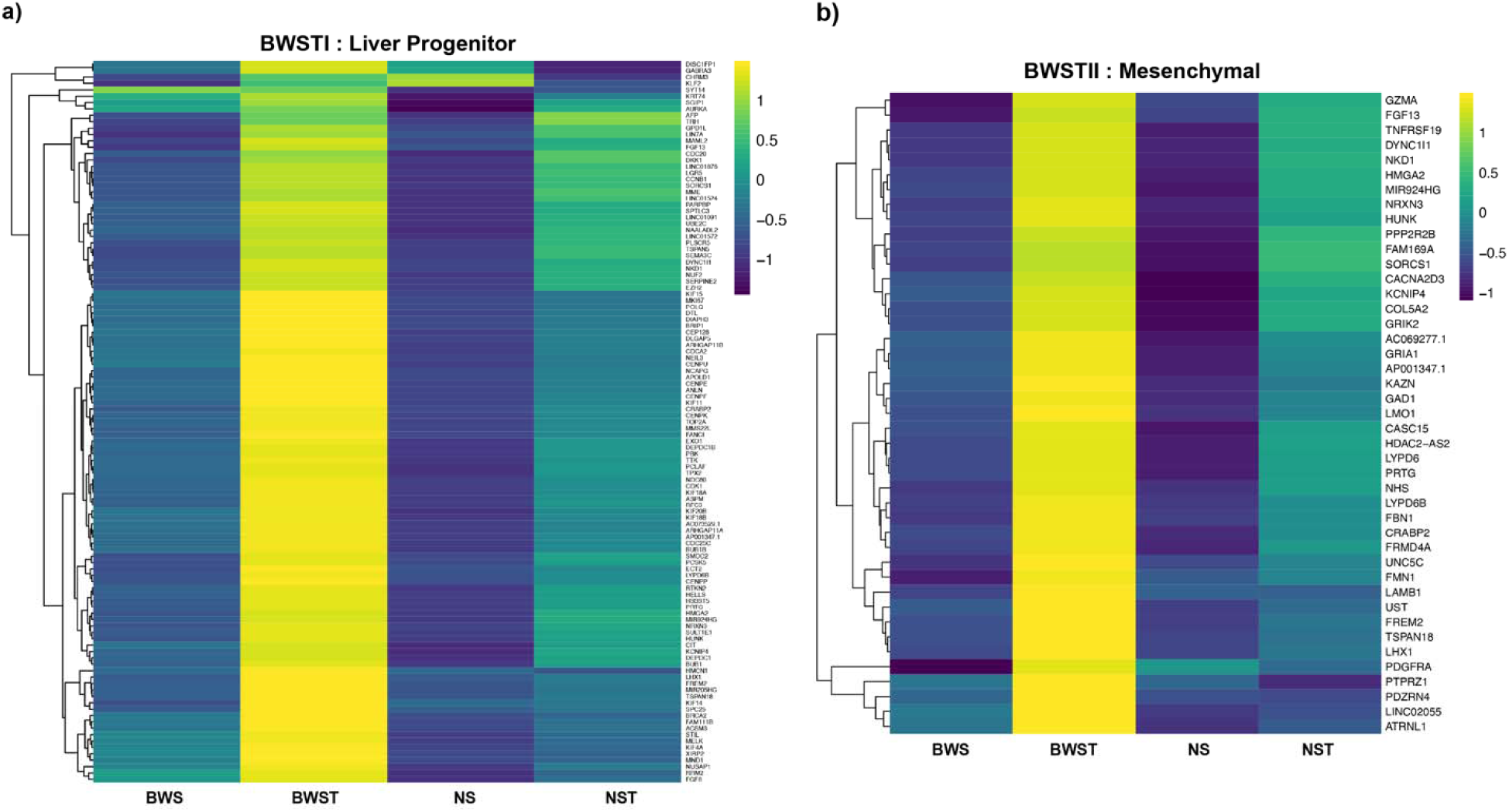
Tumor heterogeneity study reveals 2 distinct molecular signatures in BWST. Heatmaps gene expression for established BWSTI: Liver Progenitor (**a**) and BWSTII: Mesenchymal (**b**) molecular signatures in BWS tumor-adjacent liver (BWS), BWS hepatoblastoma (BWST), nonBWS tumor-adjacent liver (NS), nonBWS hepatoblastoma (NST) samples (Supplementary Data File 4).

### BWST trajectory identifies the cancer transition cell signatures

To investigate the developmental trajectory of BWS tumors compared to nonBWS, we conducted pseudotime analysis with the goal of identifying and characterizing precancerous cellular states. Given that tumor clusters predominantly grouped with hepatocytes, we selected hepatocytes and tumor clusters 1–5 from both cohorts for this analysis (**Fig. 5a, Supplementary Fig.8a**). Focusing on the distinct features of BWS hepatoblastoma, we examined the transition from hepatocytes to tumor clusters in BWS and compared these dynamics with nonBWS hepatoblastoma. Using Slingshot for trajectory inference, we identified a progression starting from normal/overgrowth/precancer/tumor-adjacent cells, advancing through a transition phase, and culminating in cancer (**Fig. 5b**). Hepatocytes were primarily localized in adjacent normal tissue and decreased in tumor tissue, while tumor clusters expanded within tumors, as anticipated (**Fig. 5a, Supplementary Fig.8a**). The pseudotime distribution of cell clusters under each condition was visualized for both cohorts (**Supplementary Fig. 8b**). In BWS, the trajectory initiated with hepatocytes, sequentially progressed through tumor clusters 1–4, and terminated at cluster 5 (**Supplementary Fig. 8b**). In contrast, the nonBWS trajectory proceeded from hepatocytes through tumor clusters 1, 3, 2, 4, and finally 5 (**Supplementary Fig. 8b**). To delineate transitioning cells—a mixed population situated between normal/tumor-adjacent and cancer tissues (56), pseudotime was divided into three bins based on cell distribution patterns: precancer (0 to 24 pseudotime), transition (24 to 34 pseudotime), and cancer (34 to max pseudotime) (**Fig. 5b**) (44, 56). We further assessed CNV changes within these intervals, observing an increase in the number of genes with non-neutral CNV states, as predicted by InferCNV’s HMM, along the pseudotime in both cohorts. This finding supported our pseudobinning approach. Cells from normal/tumor-adjacent tissue were predominantly found within the 0–24 pseudotime bin; cells from both normal/tumor-adjacent and tumor tissue populated the 24–34 bin; and cells from tumor tissue were enriched beyond 34 pseudotime (**Supplementary Fig. 9**).

**Figure 5:**
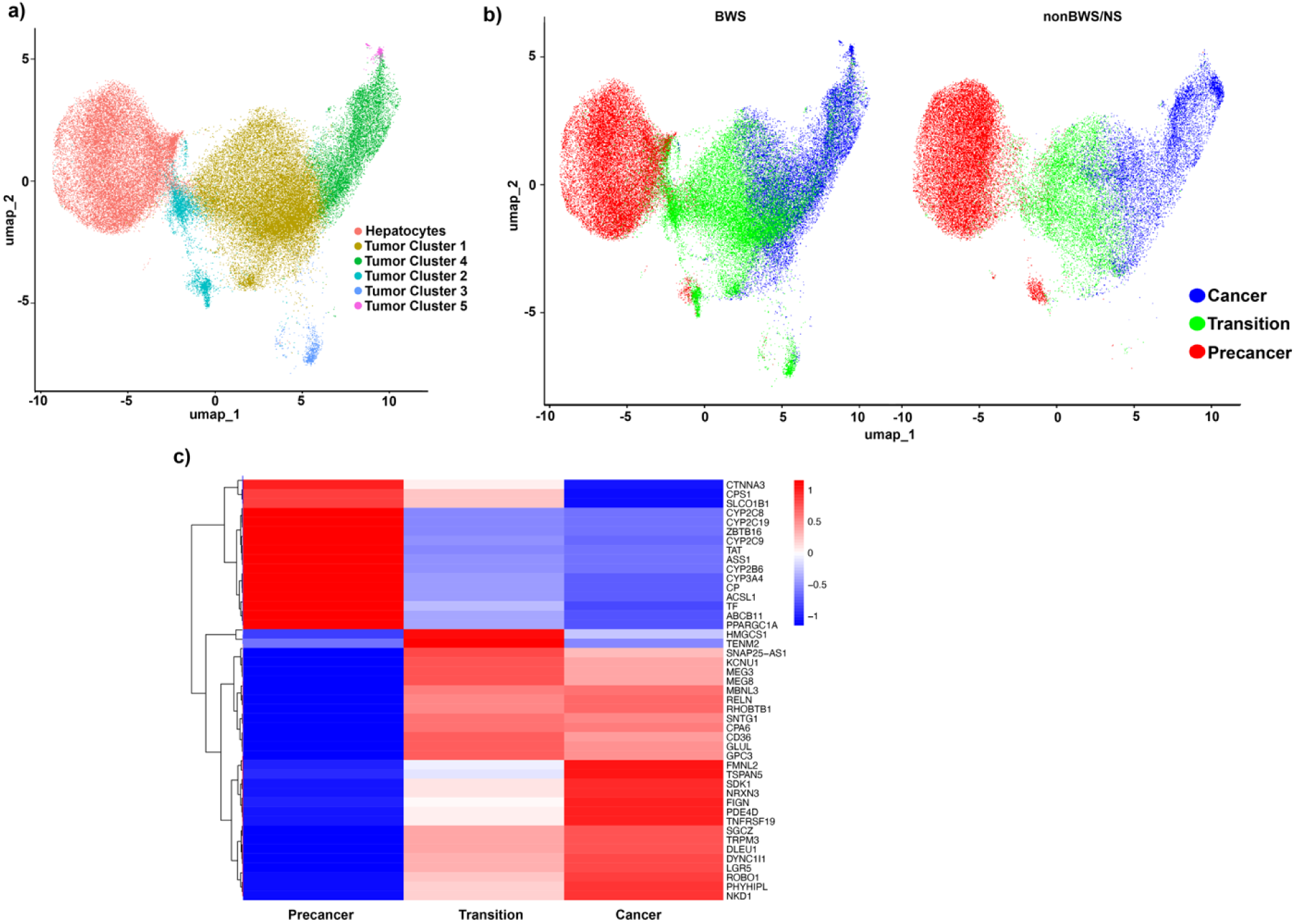
BWS hepatoblastoma trajectory analysis. (a) The UMAP of selected cell clusters used for Slingshot trajectory and pseudotime analysis. (b) The UMAP of precancer, transition and cancer state cells identified using pseudotime analysis in BWS and nonBWS samples. (c) Heatmap of expression markers expressed by precancer, transition and cancer state cells.

To characterize the cellular states identified in BWS and nonBWS tumors, we performed marker analysis (**Supplementary Data File 5**). We then plotted the average expression of approximately 15 markers across the three BWS states (**Fig. 5c)**, revealing distinct gene expression patterns reflective of functional states potentially driving tumorigenesis. Precancer cells mainly expressed liver metabolic markers *including CPS1, ASS1, CYP2B6, CYP3A4*, and *CP* (4). BWS transition cells showed expression of 14q32 locus genes (*MEG3, MEG8*) (50), the lipid metabolism regulator and ECM receptor *CD36* (57), and hepatoblastoma markers *GLUL* and *GPC3* (48, 49). BWS cancer cells expressed genes involved in Wnt signaling like *NKD1* (58) and *LGR5* (59), the TGF beta signaling molecule *TNFRSF19* (60), and ROBO1 signaling player *ROBO1* (52–54). To further elucidate the transition to malignancy, we conducted pathway analyses comparing transition versus precancer and cancer versus transition states in both cohorts (**Fig. 6a-b, Supplementary Fig.10**). In BWS transition cells, cancer-associated pathways such as KEGG_ECM_Receptor_Interaction (61) and KEGG_Gap_Junction (62) were upregulated relative to precancer cells. The cancer state exhibited broad activation of oncogenic pathways including KEGG_Wnt_Signaling, KEGG_MAPK_Signaling, KEGG_Focal_Adhesion, and KEGG_Cell_Cycle. In contrast, the nonBWS transition state was enriched for metabolic pathways including fatty acid, nitrogen, and tyrosine metabolism (**Supplementary Fig. 10a**), whereas the cancer state exhibited activation of cancer-related pathways similar to those observed in the BWS cancer state. **Supplementary Fig. 10b**). Together, these results illustrate a trajectory of pathway activation in BWS hepatoblastoma progressing from metabolically active precancer cells to transition cells with extracellular matrix remodeling, ultimately culminating in widespread oncogenic pathway activation. In nonBWS tumors, metabolic pathways predominate in the transition state before cancer pathways become activated. This analysis highlights that the BWS transition state is molecularly distinct from that of nonBWS tumors.

**Figure 6:**
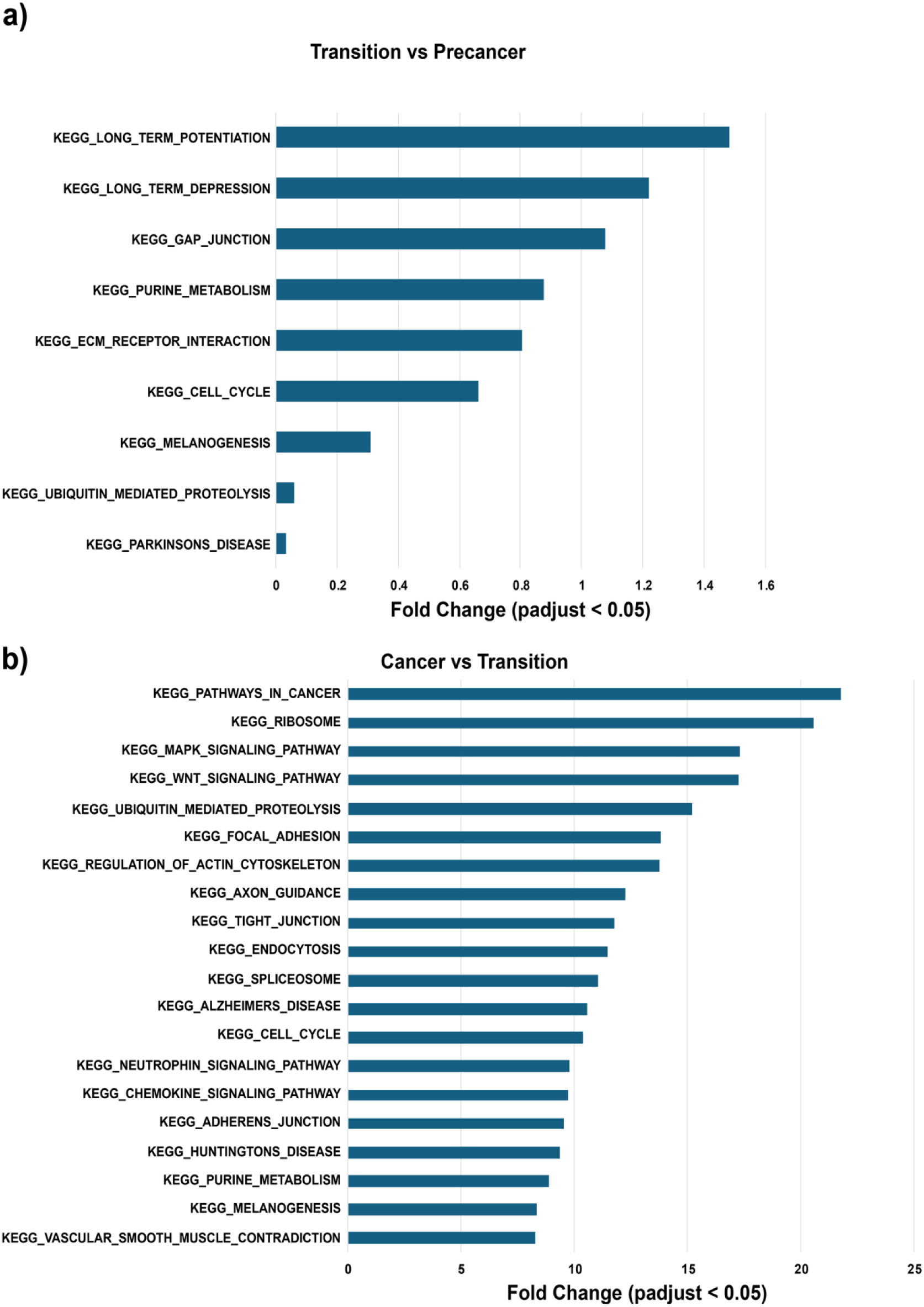
Pathway analysis for BWS hepatoblastoma trajectory analysis. (a) Pathways upregulated in transition cells relative to precancer cells. (b) Pathways upregulated in cancer cells relative to transition cells.

**Figure 7:**
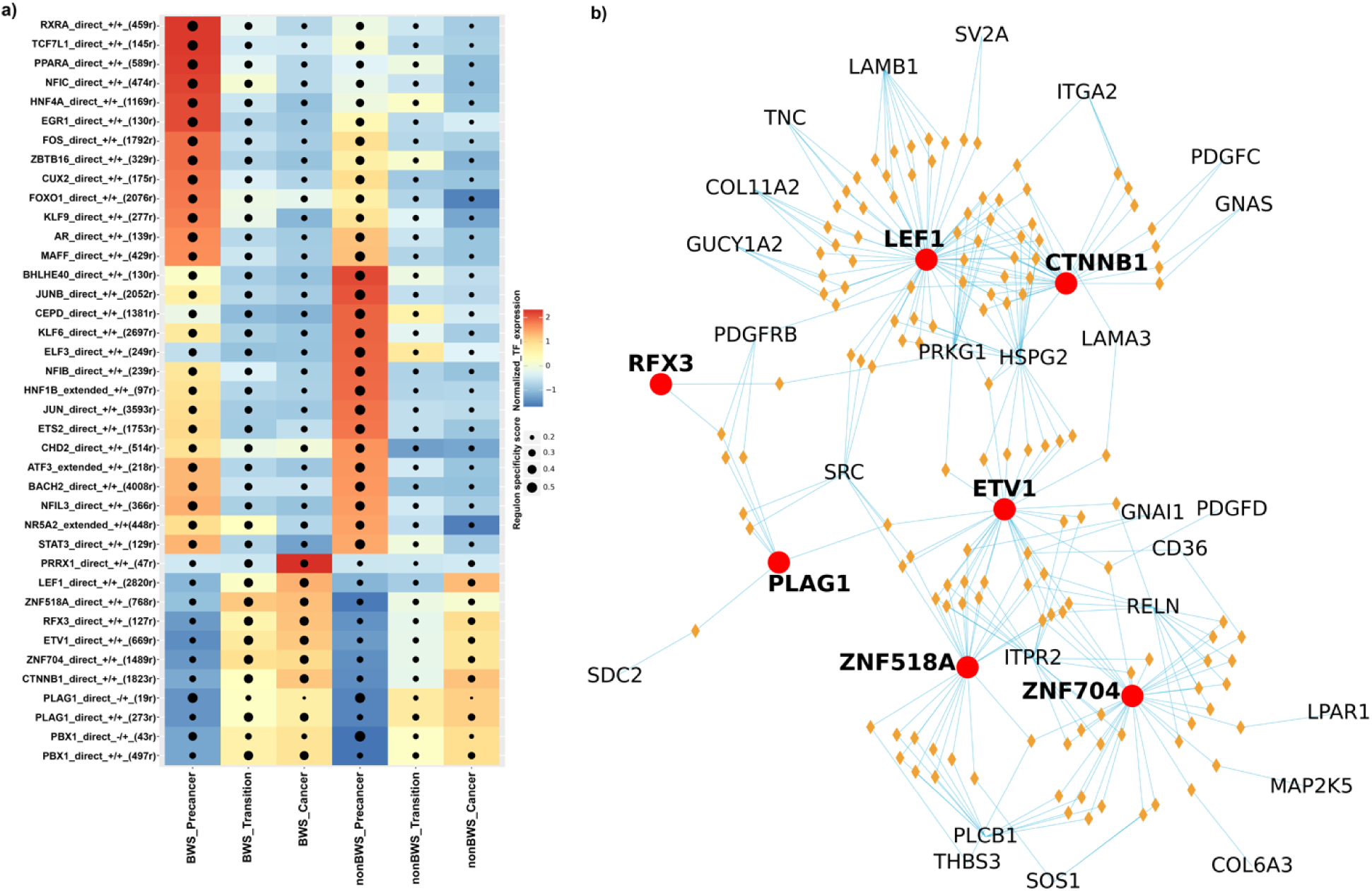
snRNA+snATAC integrated analysis using SCENIC+. **(a)** Heatmap of transcription factor expression and group specificity of the high-quality eRegulons inferred by SCENIC+ from the precancer, transition and cancer state cells of BWS and nonBWS samples. (**b**) The interaction network of transition and cancer specific eRegulons and their selected gene targets. The eRegulons include CTNNB1_direct_+/+, ETV1_direct_+/+, LEF1_direct_+/+, PLAG1_direct_+/+, RFX3_direct_+/+, ZNF518A_direct_+/+, and ZNF704_direct_+/+. The red dots represent the transcription factors of the selected eRegulons. The yellow diamonds represent the region targets of the selected eRegulons. The text labels without red dot represent the gene targets of the selected eRegulons. The blue lines represent the interactions between the transcription factors, region targets, and gene targets of the selected eRegulons.

### Integration of snRNA and snATAC datasets predict transcription factor-gene regulation network

Lastly, to investigate the transcription factor–gene regulatory networks driving the developmental trajectory of BWS hepatoblastoma in comparison to nonBWS, we integrated single-nucleus RNA-seq (snRNA-seq) and single-nucleus ATAC-seq (snATAC-seq) data from both cohorts using Single-Cell regulatory Network Inference and Clustering (SCENIC+). This analysis was conducted on hepatocytes and tumor clusters 1–5, consistent with those used in the Slingshot trajectory analysis. We compared pseudotime-defined cell states (precancer, transition, cancer) between the two cohorts. We identified 126 enhancer regulons (eRegulons) with a region-to-gene correlation coefficient (rho) > 0.55 that were differentially active across the pseudotime-defined states (**Supplementary Data File 6 – List of eregulons and transcription factors (TFs) regulated in precancer, transition and cancer states of BWS and nonBWS cohorts**). Notably, the precancer states—comprising hepatocytes in both cohorts—exhibited distinct sets of active eRegulons (**Fig. 6a**). In line with our previous observations, PPARA-driven metabolic activity was evident in BWS tumor-adjacent hepatocytes (**Fig. 6a)** (4). To better understand neoplastic transformation, we focused on eRegulons active in the transition and cancer states. We identified nine key eRegulons—PRRX1, LEF1, ZNF518A, RFX3, ETV1, ZNF704, CTNNB1, PLAG1, and PBX1, which were significantly upregulated in the transition state of BWS compared to nonBWS and maintained elevated expression in the cancer state in both cohorts (**Fig. 6a)**. To explore the functional implications of these eRegulons, we intersected the common gene set between the gene targets of these eRegulons (**Supplementary Data File 6 – List of eRegulons and transcription factors (TFs) regulated in precancer, transition and cancer states of BWS and nonBWS cohorts)** and genes involved in the Kegg_ECM_receptor_interaction, and Kegg_Gap Junction pathways – two cancer-associated pathways enriched in BWS transition cells. Using this intersection, we generated an interactome map illustrating the connections between eRegulons and their shared targets (**Fig. 6b**). The network revealed extensive interactions among the eRegulons and between their targets, highlighting a coordinated regulatory program. Specifically, CTNNB1 interacts with *ITGA2, HSPG2, PRKG1, GNAS, PDGFRB, PDGFC*; ETV1 interacts with *CD36, RELN, LAMA3, HSPG2, PDGFD, PRKG1, GNAI1, ITPR2, SRC*; LEF1 interacts with *TNC, COL11A2, LAMB1, SV2A, ITGA2, LAMA3, HSPG2, PRKG1, GUCY1A2, SRC, PDGFRB*; PLAG1 interacts with *SDC2, PDGFRB, SRC*; RFX3 interacts with *PDGFRB, PRKG1*; ZNF518A interacts with *THBS3, CD36, RELN, SOS1, PLCB1, GNAI1,SRC, ITPR2* and ZNF704 interacts with *THBS3, CD36, COL6A3, RELN, SOS1, MAP2K5,PLCB1, PDGFD, LPAR1, ITPR2*. This interactome outlines a regulatory framework in BWS transition cells that facilitates the activation of cancer-associated pathways in the subsequent cancer state.

To further investigate how this transcriptional landscape influences cell–cell communication, we applied CellChat to assess ligand–receptor interactions. We observed an enrichment of integrin-mediated signaling in BWS cancer cells compared to nonBWS, involving ligands and receptors such as *LAMC1, ITGA1/2/6, ITGB1,* and *COL4A5* (**Supplementary Fig. 11**). Notably, *LAMB1, LAMA3,* and *ITGA2*—all targets of transition-state eRegulons—were also involved in interactions during the transition phase. These results suggest that the transcriptional program in BWS transition cells establishes a molecular environment that primes cancer-state cells for enhanced signaling and malignant progression.

In summary, our integrative multi-omics analysis reveals the cellular composition, regulatory heterogeneity, and signaling architecture of BWS hepatoblastoma. We identify the transitioning cell population that bridges precancer and cancer states, characterize its distinct regulatory network, and demonstrate how this network may initiate cancer-associated signaling cascades in BWS tumors.

## Discussion

Approximately 15–20% of childhood cancers arise from an underlying cancer predisposition syndrome (63). Among these, BWS, caused by epigenetic and genetic alterations on chromosome 11p15, is the most common epigenetic cancer predisposition disorder (64, 65). Unlike many other hereditary cancer syndromes, most BWS patients lack a family history of cancer (66). BWS-associated overgrowth in the liver and kidney can lead to cancer in up to 28% of affected individuals (67). Early detection is critical, and a better understanding of the neoplastic process can improve survival and reduce long-term chemotherapy-related toxicity. Given that histology significantly influences treatment decisions in hepatoblastoma (68, 69), characterizing tumor heterogeneity is essential for developing more precise therapies, especially for advanced BWS hepatoblastoma. In this study, we delved into this research gap and performed a multiomic analysis at single-nuclei resolution to dissect the molecular landscape of BWS hepatoblastoma and its progression. Our snRNA-seq data uncovered transcriptomic features of tumor heterogeneity and delineated the developmental trajectory from tumor-adjacent tissue to tumor. We identified a population of transition cells—distinct from both normal and tumor cells—that exhibit unique gene expression profiles, open chromatin accessibility, and regulatory patterns not observed in nonBWS hepatoblastoma. Given their role in initiating tumorigenesis, these transition cells may serve as promising targets for preventive therapies.

We analyzed 147,820 nuclei from seven hepatoblastoma tumors and seven matched tumor-adjacent liver samples from four BWS and three nonBWS patients. Using our established snRNA-seq pipeline (4), we annotated all major cell types. Previously, we demonstrated that BWS tumor-adjacent tissues are metabolically hyperactive (4), and identified a BWS-specific oncogenic network via bulk transcriptomics (3). Those studies highlighted distinct molecular cues associated with 11p15 alterations. Specifically, we had identified WNT signaling as a key component of this oncogenic network (3). It has been previously reported that WNT signaling activation plays a role in tumorigenesis in patients with 11p15 alterations (24, 70), while tumor-adjacent tissues with 11p15 changes do not show expression of WNT signaling genes (24). In our current analysis, WNT signaling was enriched in both cohorts but significantly more pronounced in BWS tumors (BWST) compared to tumor-adjacent tissues. In contrast, fatty acid metabolism was more prominent in nonBWS tumors (nonBWST) and their matched tissues. These findings reinforce the critical role of WNT activation in BWS hepatoblastoma and highlight the distinct signaling patterns between BWS and nonBWS cohorts.

Despite these differences, we also identified shared oncogenic pathways. Both BWST and nonBWST samples showed enrichment in the 14q32 gene signature (50) and ROBO1 signaling, both of which are associated with poor clinical outcomes and are implicated in hepatocellular carcinoma (52–54). ROBO1 is also associated with serum alpha-fetoprotein (AFP) levels (71) - a marker routinely monitored in BWS infants for hepatoblastoma risk (6). Given that ROBO1 is detectable in serum (52), it may serve as a complementary biomarker for hepatoblastomas lacking AFP expression.

Greater than 90% of hepatoblastomas harbor mutations in *CTNNB1* (23, 72), a central WNT signaling gene found in hepatic progenitor cells that transform into tumor (14). These variants stabilize nuclear CTNNB1, forming complexes with TCF and altering the expression of WNT target genes (73). In BWS hepatoblastoma, *CTNNB1* variants—often at known hotspots—are commonly present (74). Our BWS cohort confirmed this, with 3 out of 4 BWS tumors carrying *CTNNB1* variants (3), while only one nonBWS tumor carried *CTNNB1* variant. Importantly, while 11p15 alterations are detected in BWS tumor-adjacent liver tissue, *CTNNB1* variants are restricted to tumors. This dual layer of molecular complexity, involving both epigenetic (11p15) and somatic (*CTNNB1*) alterations, distinguishes BWS hepatoblastoma from its nonBWS counterpart. To further characterize this heterogeneity, we analyzed hepatoblastoma molecular markers across BWS and nonBWS tumors. BWS tumors expressed gene signatures indicative of embryonal, liver progenitor, and mesenchymal states, consistent with prior reports (13–15). Feka *et al* described a BWS hepatoblastoma with mixed epithelial-mesenchymal histology (13), while other studies reported embryonal hepatoblastomas lacking mature hepatic markers but enriched in WNT signaling (14, 15).

Using pseudotime analysis (25, 26, 28, 29, 44, 56), we modeled the trajectory from normal to cancerous liver in BWS. We identified three distinct states: precancer, transition, and cancer. The precancer state comprised metabolically active hepatocytes, enriched in liver metabolic genes. In contrast, the cancer state was defined by upregulation of key oncogenic pathways, including WNT signaling components such as *NKD1* (58), *LGR5* (59), *TNFRSF19* (60) and *ROBO1*. The transition state exhibited a unique gene expression profile, marked by *MEG3*, *MEG8*, *HMGCS1*, and *CD36*. Notably, *MEG3* and *MEG8* have opposing effects in hepatocellular carcinoma—*MEG3* is tumor-suppressive (75), while *MEG8* is oncogenic (76).Their co-expression may influence clinical outcomes in BWS hepatoblastoma.

*HMGCS1*, involved in cholesterol synthesis, is upregulated in the BWS transition state and is known to respond to *PPARA* activation (77). This aligns with our previous finding that *PPARA* is overexpressed in BWS liver (4). Cholesterol accumulation has been linked to malignancy via collagen-integrin signaling (78), and our data showed increased collagen and integrin signaling in the BWS cancer state. *CD36*, another transition marker, regulates lipid metabolism and is an extracellular matrix (ECM) receptor known to foster immunosuppressive tumor microenvironments (57). Pathway enrichment analyses confirmed that ECM receptor interaction and gap junction pathways were hallmarks of the BWS transition state, while WNT signaling was dominant in the cancer state. Considered as a temporal progression model, the BWS precancer state is metabolically active. Through mechanisms that are not yet fully understood, it initiates the expression of ECM-related pathways in BWS transition cells.

We further integrated snRNA-seq with snATAC-seq using SCENIC+ to identify key transcriptional regulators across the three cell states. The BWS precancer state featured eRegulons including PPARA, RXRA, and TCF7L1, which drive its metabolic activity. In contrast, the nonBWS precancer state featured different regulators (e.g., BHLHE40, JUNB, CEBPD, KLF6) involved in general liver metabolism (79–82). The transition and cancer states in BWS showed elevated activity of several transcription factors: PRRX1, LEF1, ZNF518A, RFX3, ETV1, ZNF704, CTNNB1, PLAG1, and PBX1. Network analysis revealed that many BWS transition state eRegulons regulate ECM-related genes such as *ITGA2*, *LAMB1*, and *LAMA3*, which are further upregulated in cancer cells. For instance, LAMA3 promotes epithelial-mesenchymal transition (EMT) in cholangiocarcinoma (83), while ETV1 and PBX1—upregulated in BWS—are associated with EMT and metastasis in hepatocellular carcinoma (84) (85). PLAG1, another transition state regulator, activates *IGF2*, a gene frequently upregulated in BWS (86) (3).

Our working hypothesis posits that *PPARA* overexpression in the BWS precancer state induces downstream targets, including *CD36* (87). This leads to metabolic alterations, elevated oxidative stress (4), and increased circulating free fatty acids (88). These, in turn, activate *CD36*, which has been shown to promote EMT via WNT and TGF-β pathways (89, 90). Additionally, the increased reactive oxygen species (ROS) may contribute to the acquisition of somatic *CTNNB1* variants, driving tumor formation. Thus, targeting key signaling pathways in the transition may help in the prevention of hepatoblastoma in BWS.

Although our study was limited by sample size, it provides an in-depth characterization of tumor specimens, demonstrating how this level of analysis can offer valuable insights into tumor biology. In conclusion, in this study, we characterized the epigenetic and transcriptional landscape of BWS-associated hepatoblastoma and its matched tumor-adjacent liver tissue, providing a detailed comparison with nonBWS samples. Our analysis revealed distinct tumor heterogeneity associated with BWS and identified key molecular features that differentiate BWS hepatoblastoma from its nonBWS counterpart. Importantly, we delineated a stepwise progression from normal liver to cancer in BWS, uncovering specific pathways involved in this transition. We identified a unique population of transition cells with mixed normal and tumor-like genetic features, which appear to mediate the progression from a precancerous to a malignant state. These cells may represent a critical target for early intervention or prophylactic therapy aimed at preventing tumor initiation in high-risk BWS patients. Further investigation of this transitional cell population and its regulatory mechanisms could pave the way for novel therapeutic strategies in cancer prevention and management for BWS.

## Acknowledgements

We would first and foremost like to thank the patients and families who are members of the BWS Registry and provided their tissue samples for research purposes. We thank CHOP Center for Applied Genomics, for performing the sequencing. We thank Dr. Kate Creasy and Amrith Rodrigues from Dr. Daniel Rader’s lab at the University of Pennsylvania for their assistance with optimizing the single-nuclei extraction from samples. We thank Dr. Zhen Miao from Junhyong Kim’s lab of University of Pennsylvania for helping us setting up PICsnATAC software for our snATAC study. We also thank Dr. Rebecca Linn from the Division of Anatomic Pathology at CHOP for coordinating the nonBWS liver samples used in this study; we would also like to respectfully acknowledge the individuals from whom these samples were derived.

## Authors’ contributions

S.N., E.D.T., and J.M.K. designed the study. S.N. and R.D.P. performed experiments. S.N., E.D.T., R.D.P., S.P.M., Y.Z., and M.X. analyzed the data. K.M.B provided materials as part of the CHOP Center for Childhood Cancer Research biobank and clinical data. S.N., E.D.T., and J.M.K. wrote the manuscript.

## Ethics approval and consent to participate

All the samples and clinical information from patients with BWS were collected through the BWS Registry established at the Children’s Hospital of Philadelphia under an approved institutional review board protocol (IRB 13-010658). We confirm that all research was performed in accordance with the relevant guidelines/regulations of the Children’s Hospital of Philadelphia under an approved institutional review board protocol (IRB 13-010658). The study protocols were in compliance with the revised Declaration of Helsinki. The liver/tumor adjacent and hepatoblastoma samples of nonBWS patients were collected under the protocol (IRB 21-018450_AM13). Informed consent was obtained for the collection of clinical information and the samples from all partcipants comprising this study. The limited clinical information regarding the nonBWS samples was extracted from the electronic clinical record through an honest broker.

## Consent for publication

Consent was obtained from all patients and/or legal guardians to collect longitudinal clinical information to publish the findings.

## Data availability

Raw sequencing data from patient samples (snRNA-seq) were deposited in dbGAP with accession number phs002614.v2.p1. This study did not generate any unique code. All software tools used in this study are publicly available. The authors declare that all R scripts supporting the findings of this study are available from the corresponding author upon reasonable request. The graphical abstract was generated using Biorender with agreement number BK28BMPMHT.

## Supplementary Figures and Tables

**Supplementary Figure 1:**
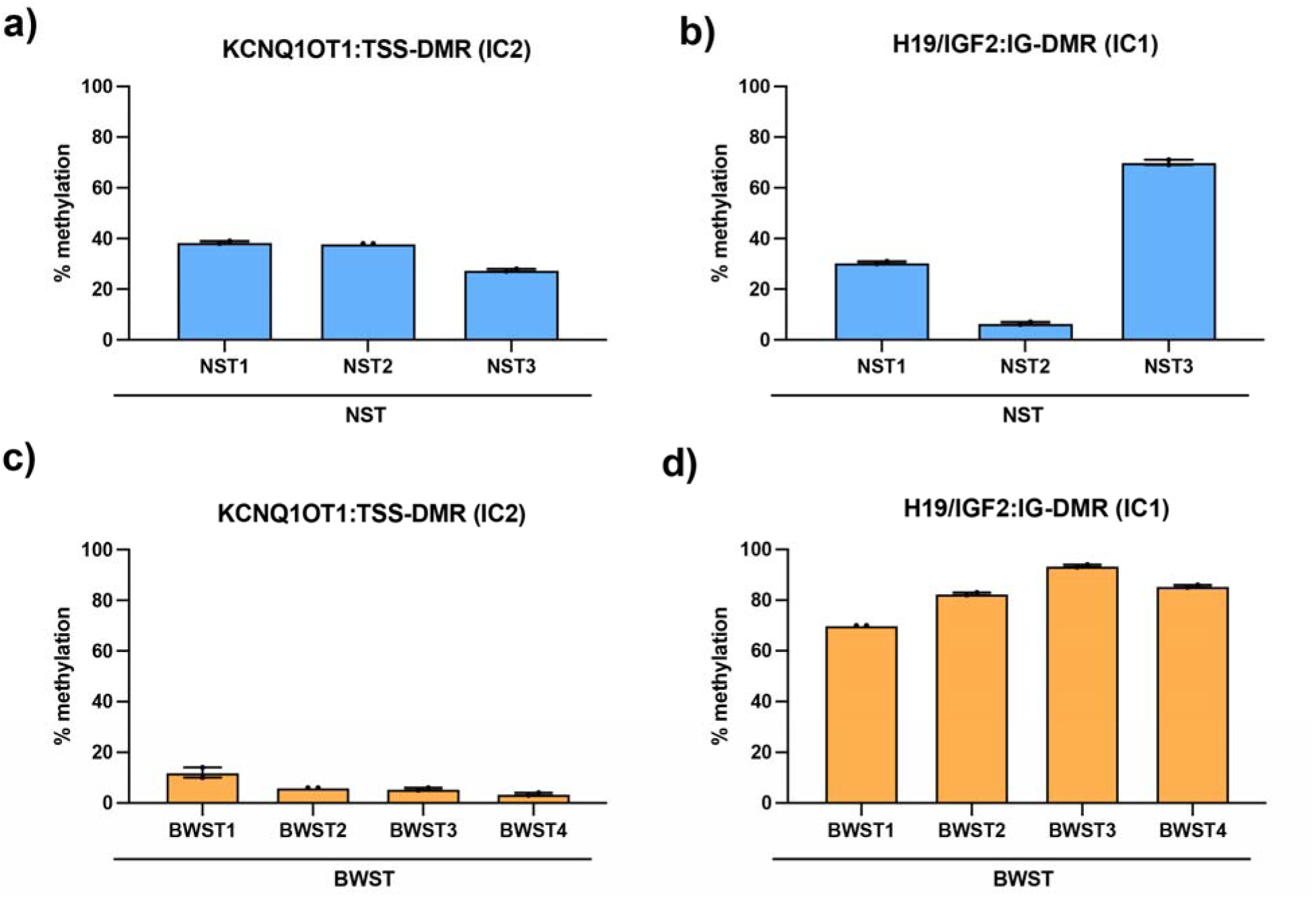
Methylation profiles of hepatoblastoma samples. Methylation profiles of nonBWS (NS) tumor/hepatoblastoma samples were assessed for IC2 (KCNQ1OT1:TSS-DMR; **a**) and IC1 (H19/IGF2:IG-DMR; **b**). Similarly, BWS tumor/hepatoblastoma samples were assessed for IC2 (**c**) and IC1 (**d**). These methylation profiles were measured by bisulfite pyrosequencing, as described in methods. Displayed are mean ± SEM from 2 replicates. BWST1 corresponds to Patient 7, BWST2 corresponds to Patient 6, BWST3 corresponds to Patient 3 and NWST4 corresponds to Patient 2 from published BWS bulk transcriptomics study (1). Supplementary table 1 shows the methylation profile for nontumor tissue that we have previously reported.

**Supplementary Figure 2:**
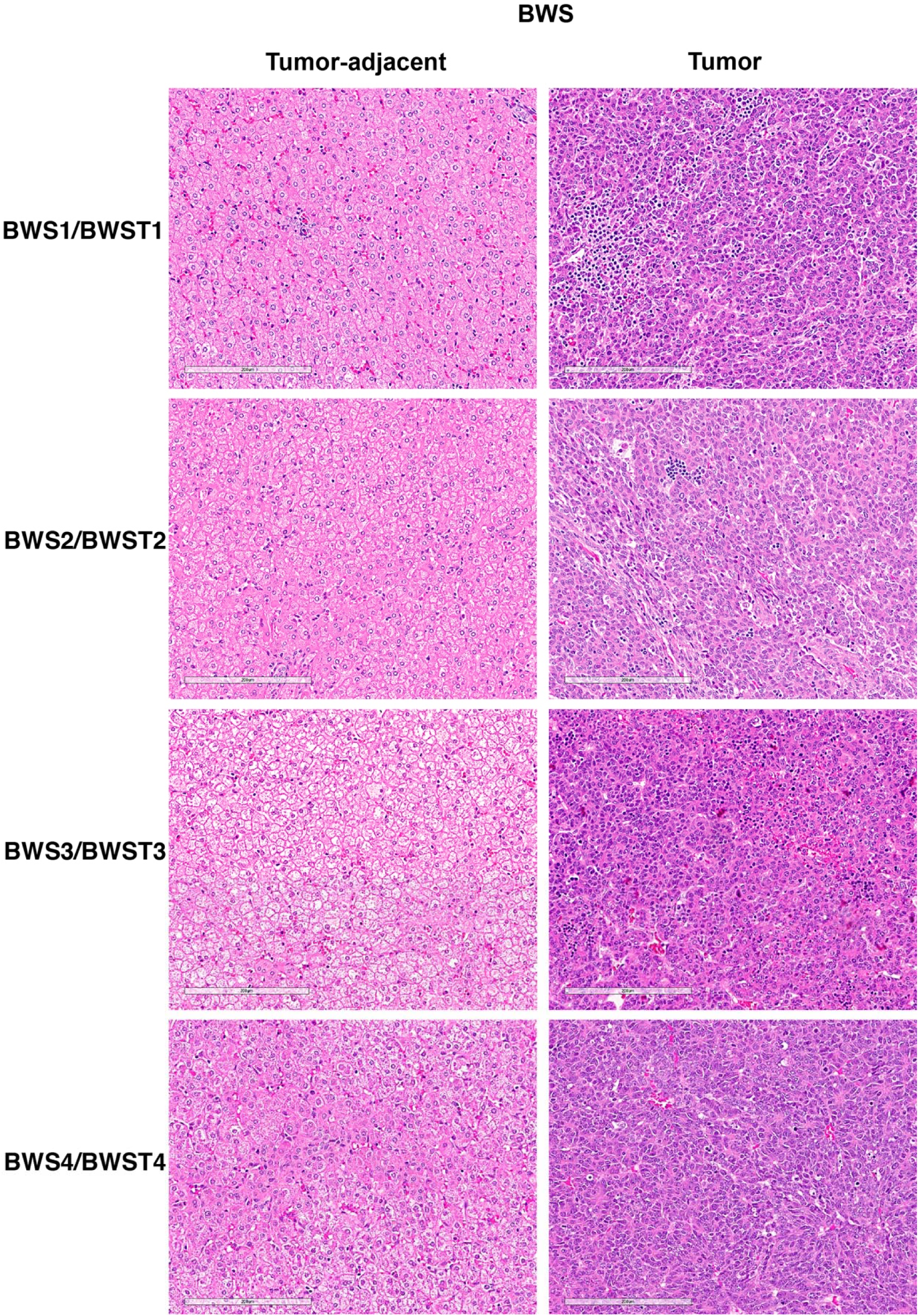
Assessment of BWS normal liver and tumor histology. Representative liver sections from BWS tumor adjacent liver and tumor were stained with hematoxylin and eosin. Scale bar: 200 μm.

**Supplementary Figure 3:**
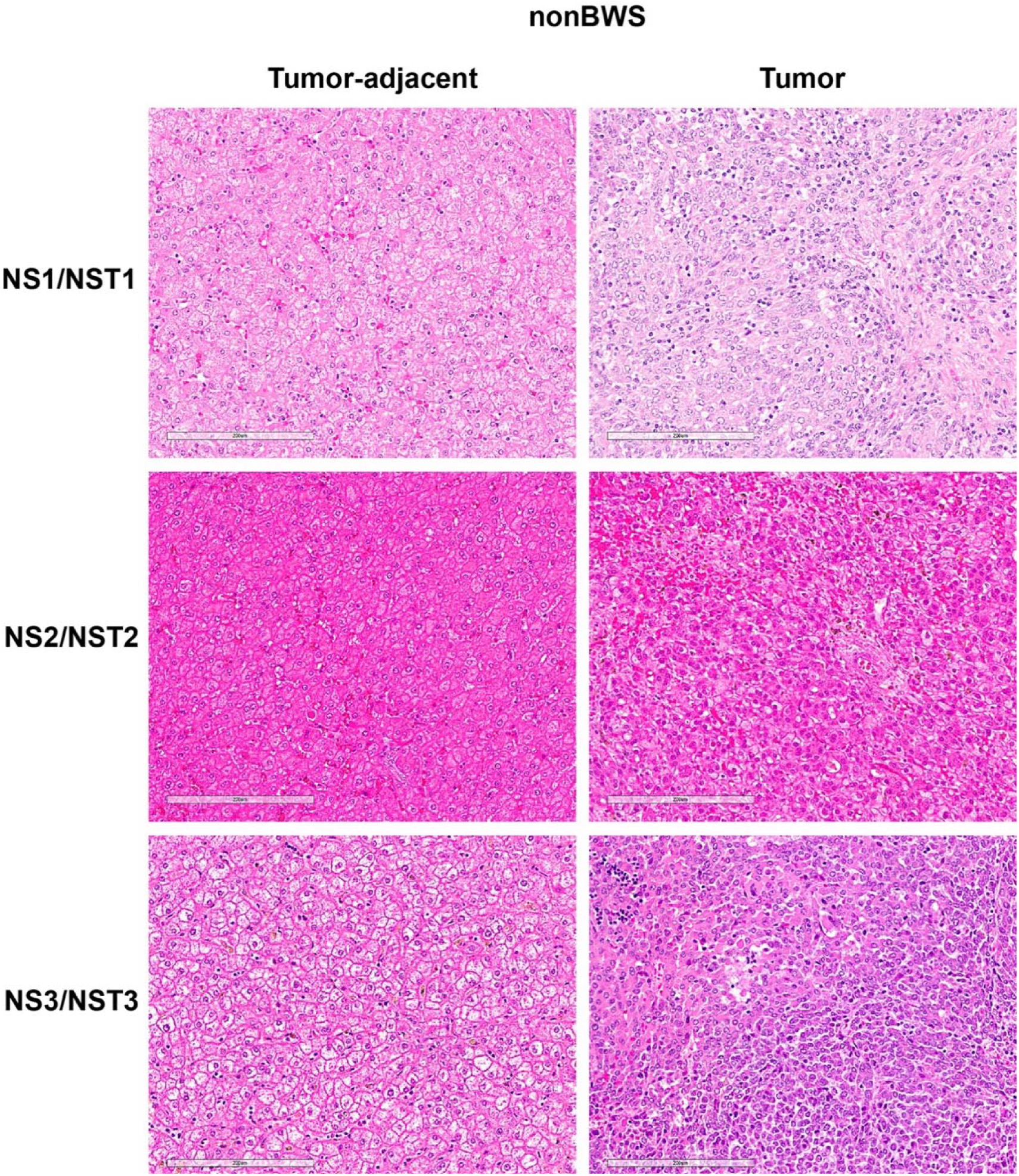
Assessment of nonBWS normal liver and tumor histology. Representative liver sections from nonBWS tumor-adjacent liver and tumor were stained with hematoxylin and eosin. Scale bar: 200 μm.

**Supplementary Figure 4:**
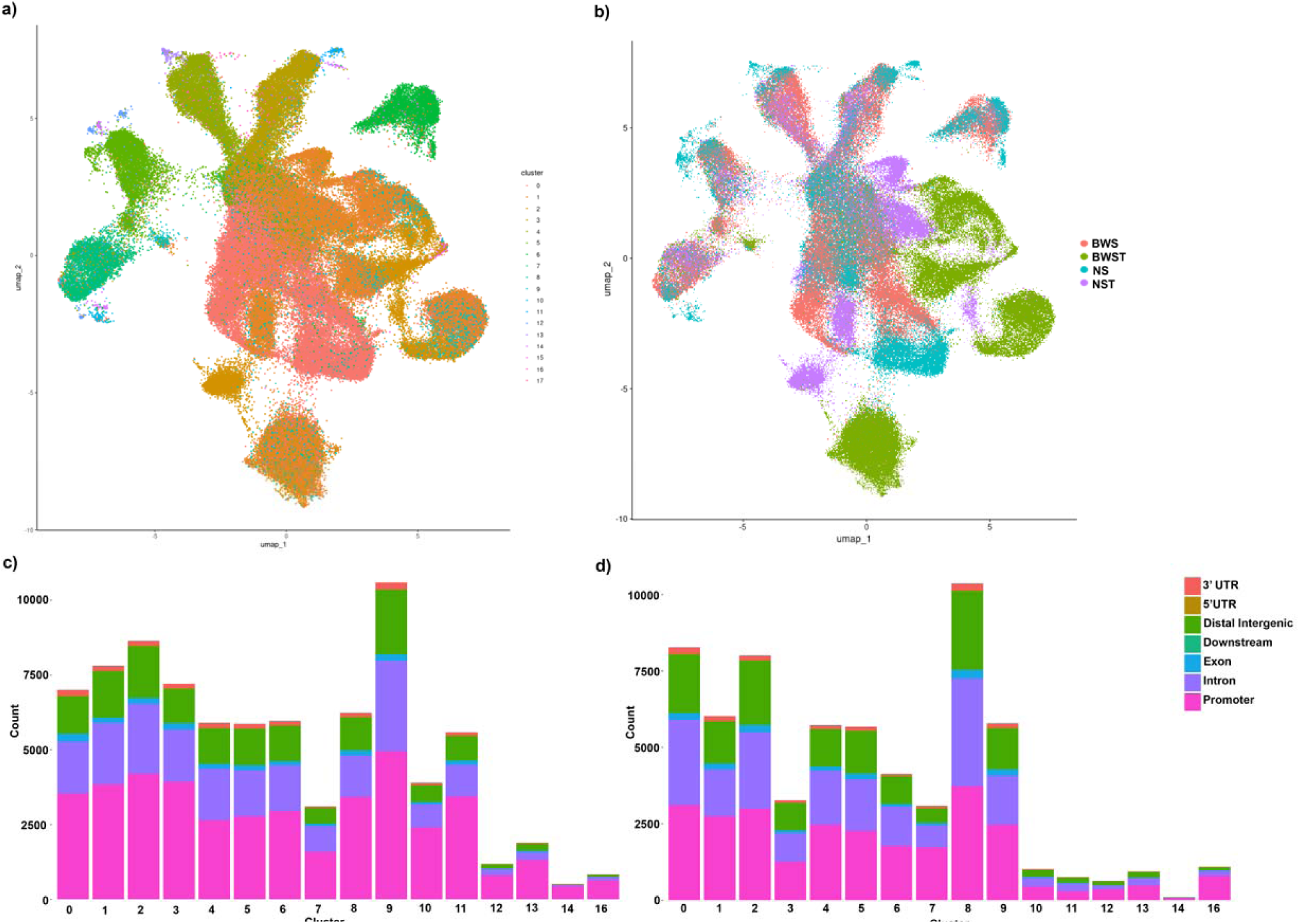
Chromatin accessibility profile of BWS (tumor-adjacent and hepatoblastoma/tumor) and nonBWS (tumor-adjacent and hepatoblastoma/tumor) samples. The UMAP of 18 cell clusters representing the BWS (tumor-adjacent and hepatoblastoma/tumor) and nonBWS (tumor-adjacent and hepatoblastoma/tumor) samples (a). The UMAP of tumor-adjacent and tumor distribution of BWS and nonBWS samples (b). Bar plot of peaks enriched in UTR, promoter, distal, intronic and exonic regions of BWS (c) and NS/nonBWS (d) samples.

**Supplementary Figure 5:**
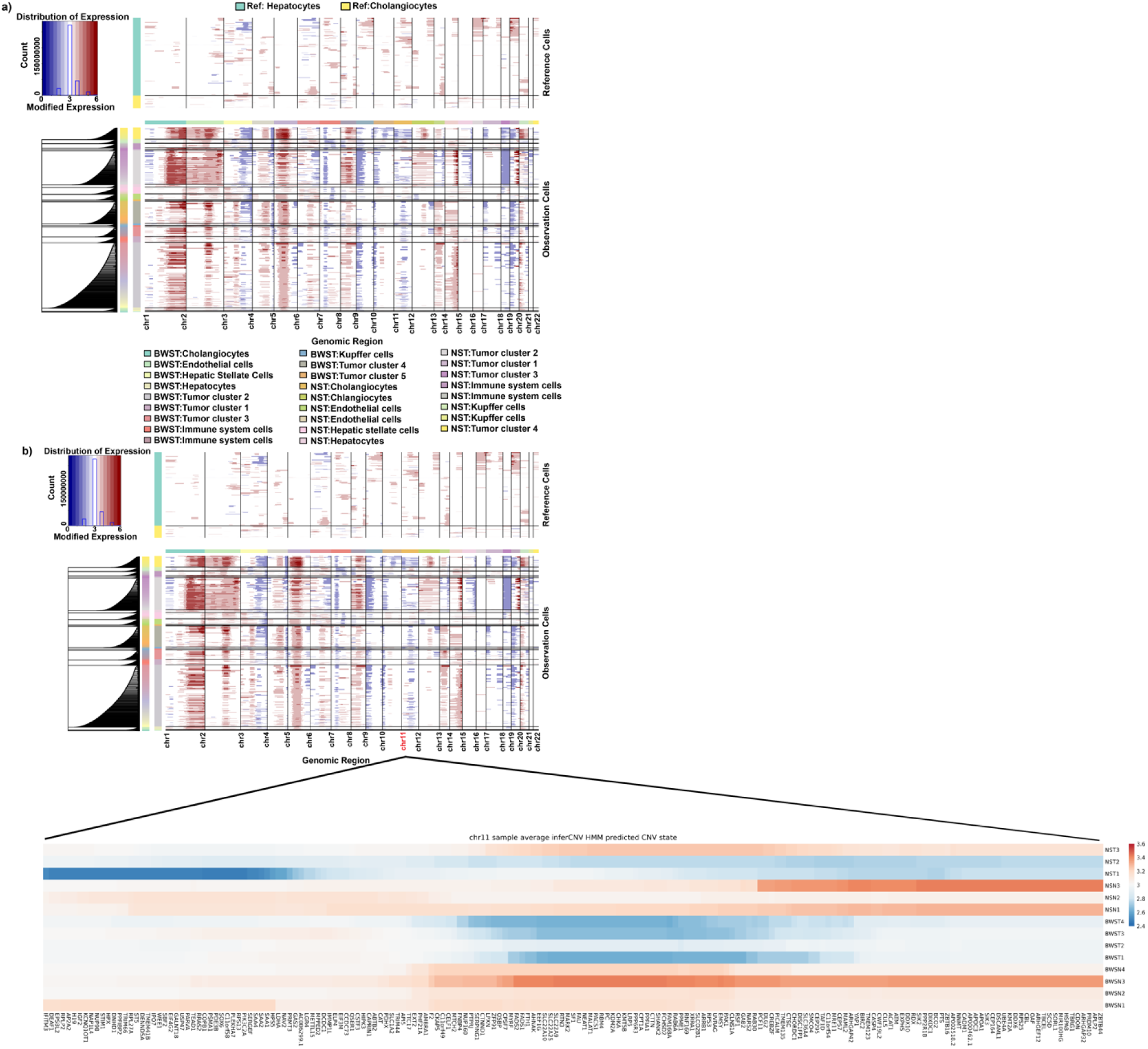
Copy number alteration profile. (a) Estimated copy-number alteration profile of all liver cell and tumor clusters from both BWS and nonBWS (NS) samples using the hepatocytes and cholangiocytes cells from non-tumor samples (BWS and NS) as reference. Estimated copy numbers are shown in blue (deletion) and red (amplification) color bars. (b) Heatmap displaying estimated copy-number alteration profile of chromosome 11 for all samples in study. The gene copy-number changes shown in the heatmap are the average InferCNV HMM predicted states of all the cells in a given sample. The color bar gives the Sample average InferCNV HMM predicted CNV state.

**Supplementary Figure 6:**
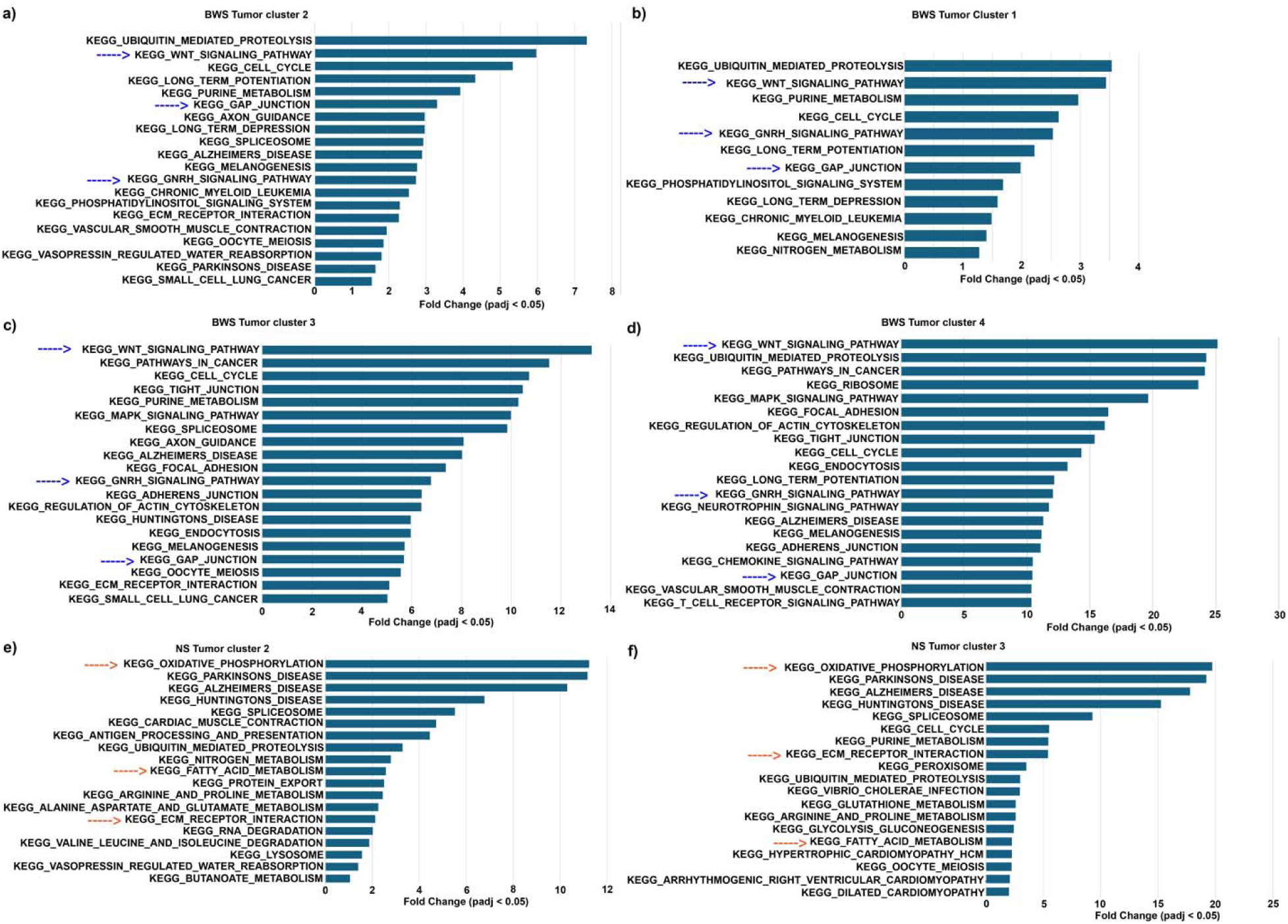
Pathway enrichment analysis in tumor clusters. Pathways enriched in BWS tumor cluster 2 (a), BWS tumor cluster 1 (b), BWS tumor cluster 3 (c), BWS tumor cluster 4 (d), nonBWS (NS) tumor cluster 2 (e), nonBWS (NS) tumor cluster 3 (f) **Supplementary Data File 3**. nonBWS (NS) tumor cluster 1 showed enrichment of Kegg_Ribosome and Kegg_Steroid_biosynthesis pathway. nonBWS (NS) tumor cluster 4 showed enrichment of KEGG_TIGHT_JUNCTION, KEGG_NON_SMALL_CELL_LUNG_CANCER, KEGG_PHOSPHATIDYLINOSITOL_SIGNALING_SYSTEM, KEGG_ADHERENS_JUNCTION, KEGG_VEGF_SIGNALING_PATHWAY (**Supplementary Data File 3**).The blue arrows indicate the common pathways in BWS tumor clusters while orange arrows indicate the common pathways in nonBWS tumor clusters.

**Supplementary Figure 7:**
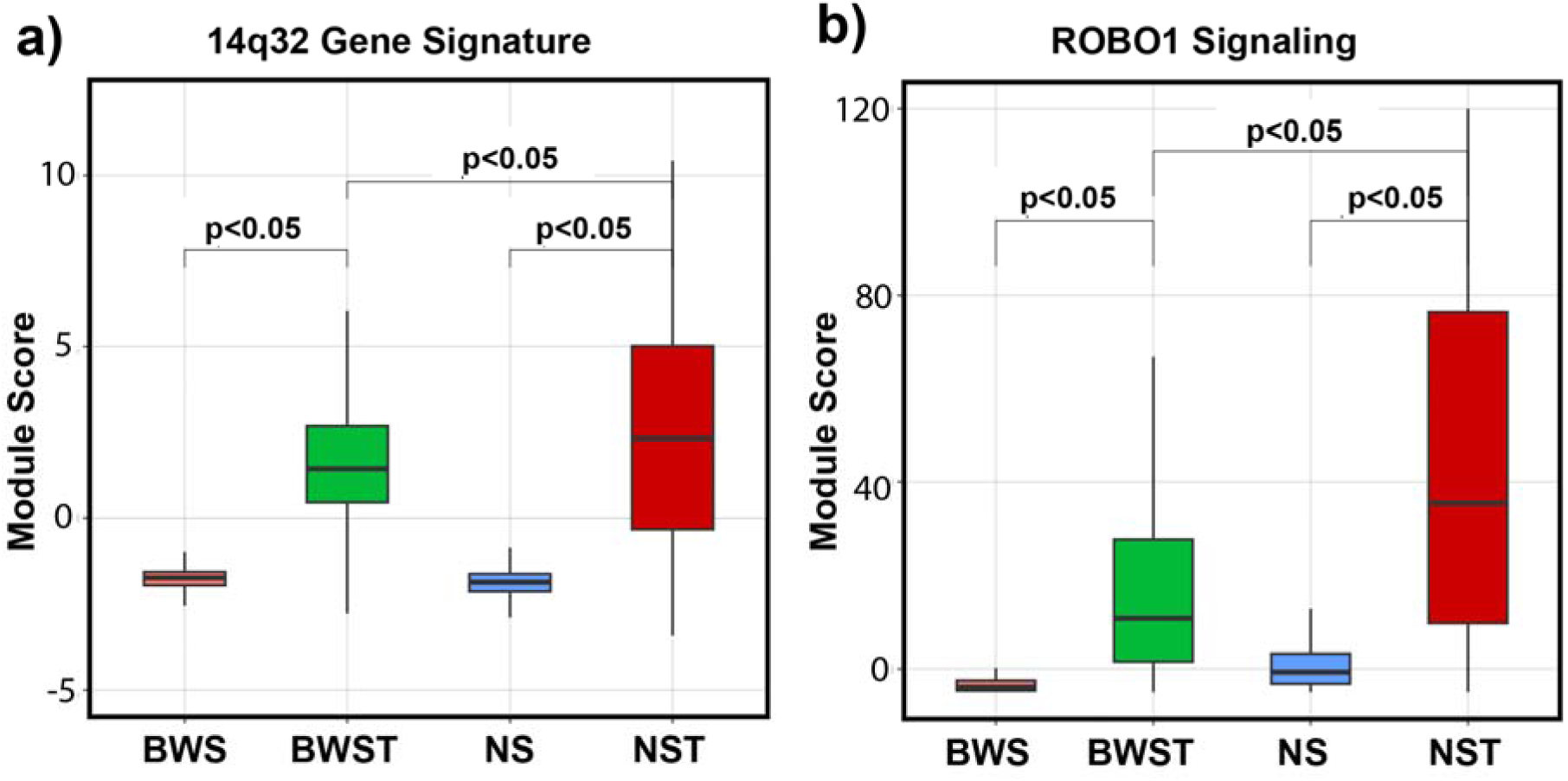
Examination of previously reported liver tumor markers in BWS and nonBWS hepatoblastoma. Box plots showing module score of the 14q32 gene signature (**a**) and ROBO1 signaling (**b**) in liver and hepatoblastoma samples of BWS and nonBWS (NS) cohorts.

**Supplementary Figure 8:**
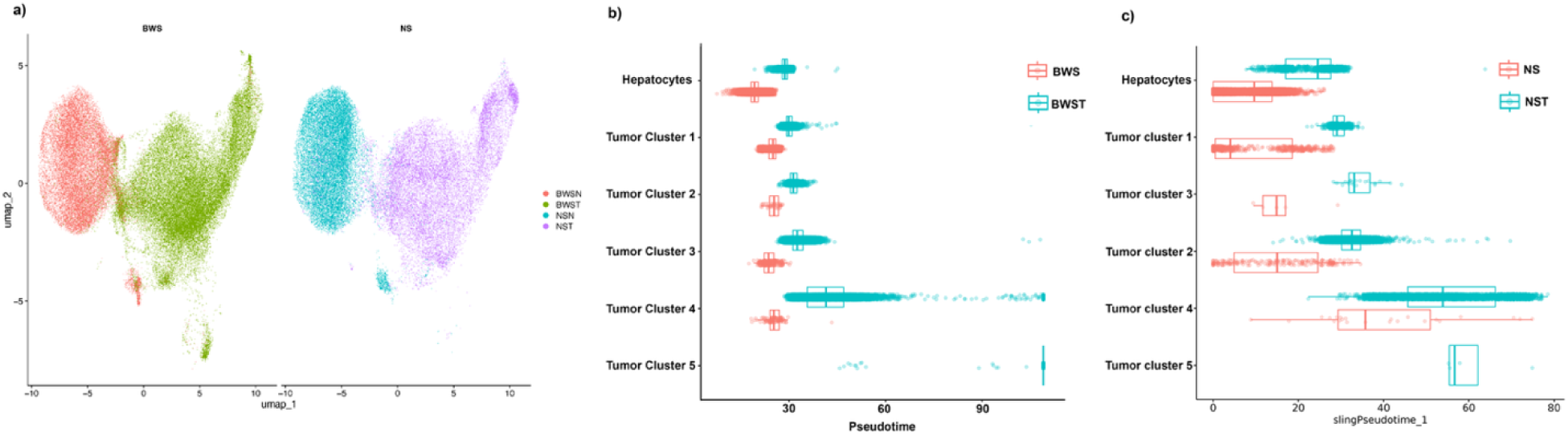
Normal and tumor cell distribution in BWS and nonBWS/NS samples and nonBWS trajectory analysis. (**a**) UMAP showing distribution of normal and tumor cell distribution in BWS and nonBWS samples. (**b**) Boxplot showing the distribution of pseudotime values for each selected clusters partitioned by the normal/nontumor or tumor condition in BWS and nonBWS cohorts. (**c**) The UMAP of precancer, transition and cancer state cells identified using pseudotime analysis in BWS and nonBWS samples.

**Supplementary Figure 9:**
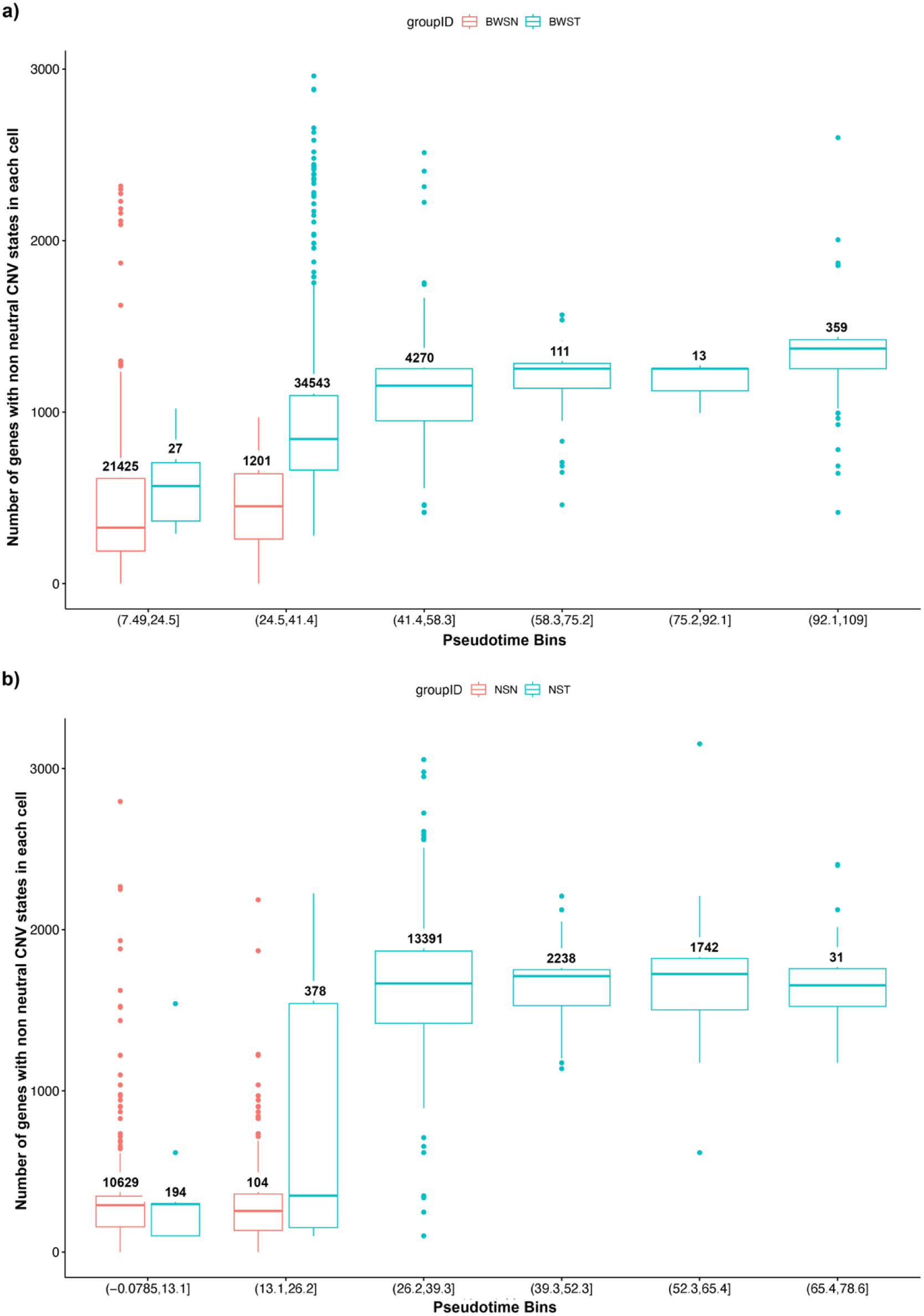
Virtual copy number profile changes with pseudotime. Boxplot of the number of genes with non-neutral InferCNV HMM predicted CNV states in each cell, with the cells in BWS (**a**) and nonBWS (**b**) tumor-adjacent and tumor tissues grouped by pseudotime bins.

**Supplementary Figure 10:**
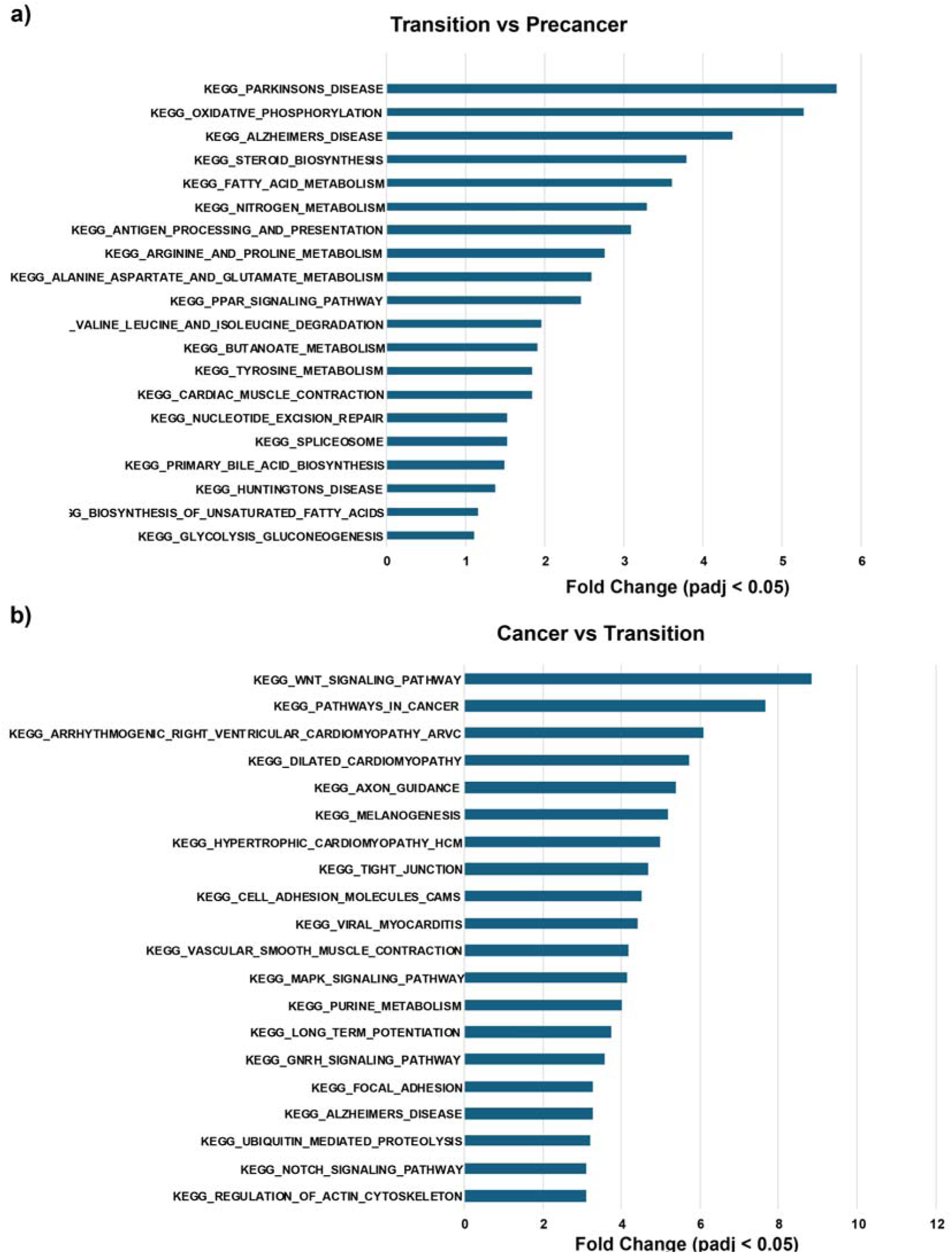
Pathway analysis for nonBWS pseudotime states. Pathways upregulated in nonBWS transition cells relative to nonBWS precancer cells (**a**). Pathways upregulated in nonBWS cancer cells relative to nonBWS transition cells (**b**).

**Supplementary Figure 11:**
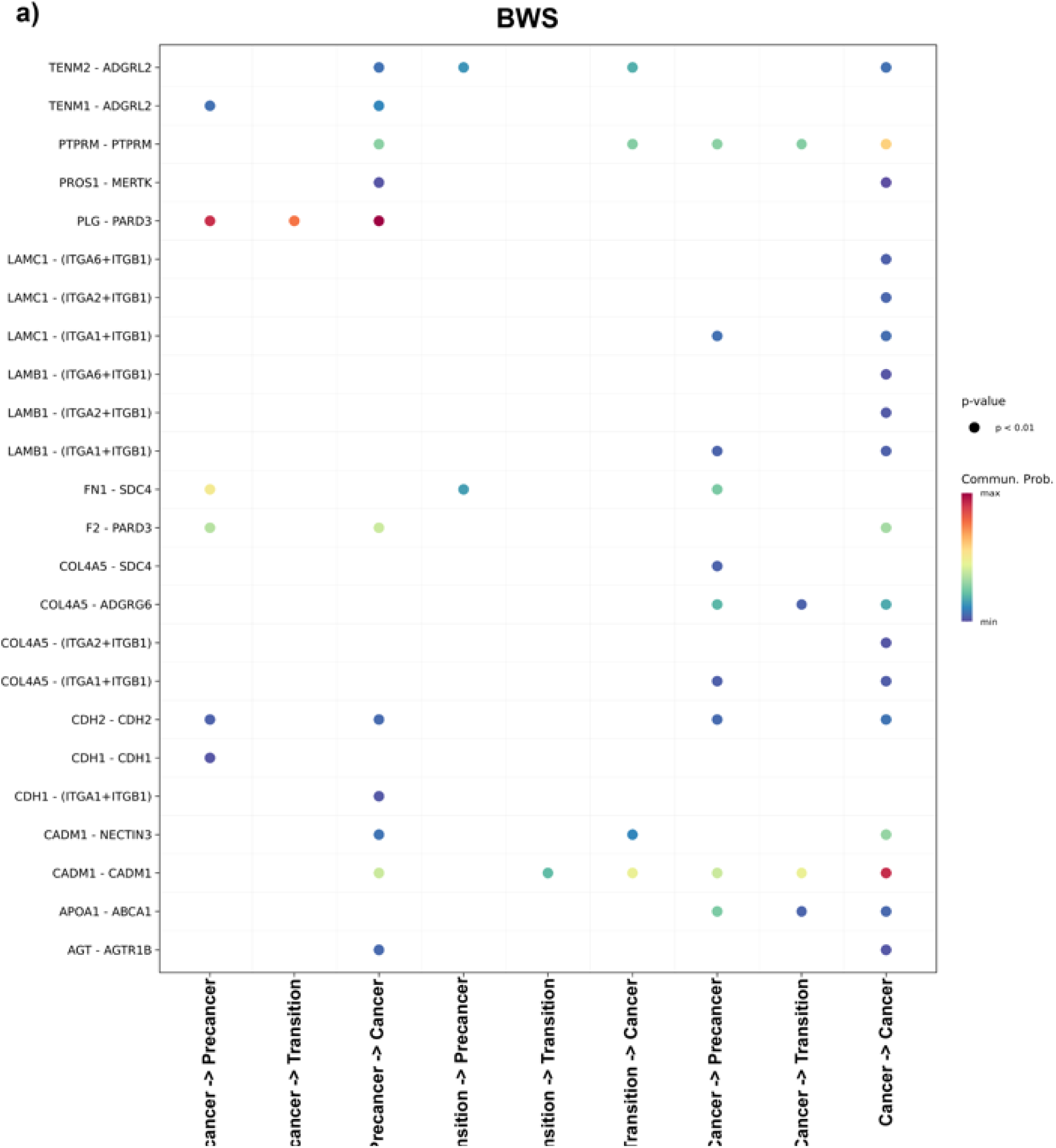
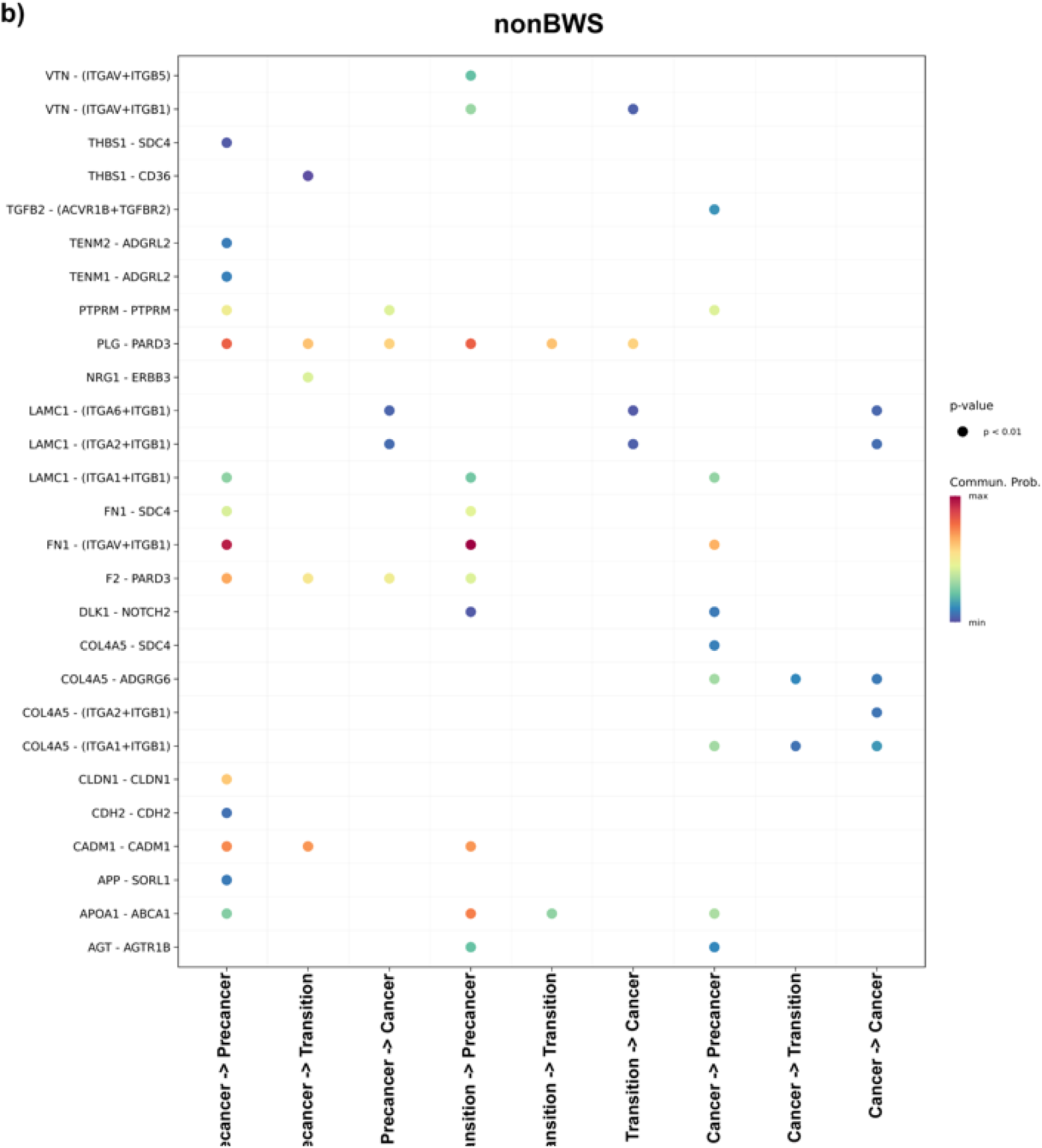
Ligand-receptor interactions study. Cellchat inferred ligand-eceptor interaction probabilities between precancer (0–24), transition (24–34) and cancer (34–110) state cells predicted in BWS (a) and nonBWS (b) samples. The receptor-ligand pairs are shown on Y axis while X axis shows the source-target cells pair.

**Supplementary Table 1:** % Methylation at IC1 (H19/IGF2:IG-DMR) and IC2 (KCNQ1OT1:TSS-DMR) for tumor adjacent/nontumor tissues in study. Data points adapted from (2).

| Sample Name | % methylation at IC1<br>(H19/IGF2:IG-DMR) | % methylation at IC2<br>(KCNQ1OT1:TSS-DMR) |
| --- | --- | --- |
| BWS1 | 20.30 | 24 |
| BWS2 | 30.08 | 17 |
| BWS3 | 34.87 | 20.5 |
| BWS4 | 19.12 | 4 |
| NS1 | 35.5 | 38 |
| NS2 | 9.5 | 37.5 |
| NS3 | 30.5 | 37.5 |

**Supplementary Table 2:**
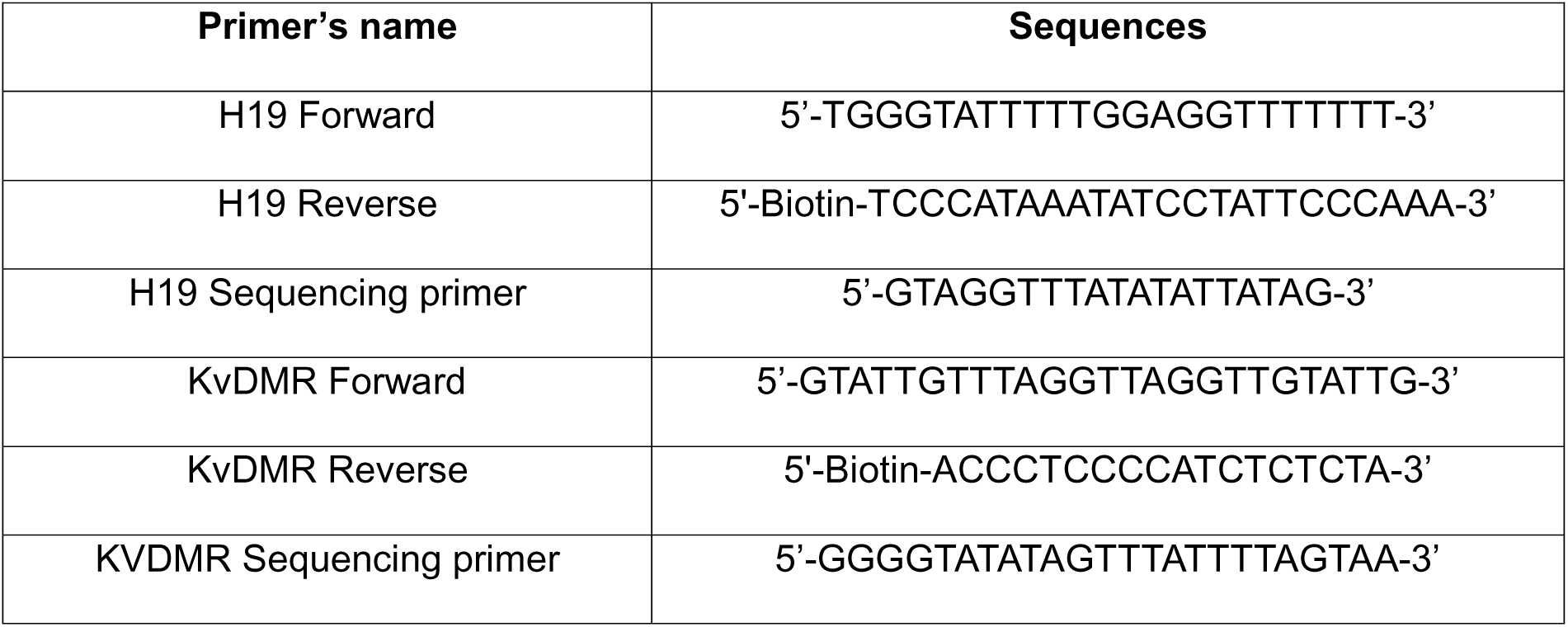
Primer sequences used for pyrosequencing.

**Supplementary Table 3:** Clinical information on the cohort used in multiomics study (2).

| Sample ID | Cohort | BWS subtype in blood | % Methyl (IC2) | % Methyl (IC1) | % Mosaicism in blood | Sex | Age at Surgery | Tumor histology | CTNNB1 mutation in tumor |
| --- | --- | --- | --- | --- | --- | --- | --- | --- | --- |
| BWS1 | BWS | IC2 LOM | 27.12 | 50.41 | 45.76 | F | 5 months | Mixed fetal and embryonal | Yes |
| BWS2 | BWS | pUPD11 | 36.99 | 61.33 | 24.34 | M | 13 months | Mixed fetal and embryonal | Yes |
| BWS3 | BWS | pUPD11 | 16.87 | 80.27 | 63.4 | F | 14 months | Mixed fetal and embryonal | Yes |
| BWS4 | BWS | IC2 LOM | 00.05 | 48.16 | 99.9 | M | 2 months | Embryonal with scattered mesenchymal foci | No |
| nonBWS1 | nonBWS |  |  |  |  | M | 2 months | Epithelial and mesenchymal | No |
| nonBWS2 | nonBWS |  |  |  |  | F | 1 month | Embryonal | No |
| nonBWS3 | nonBWS |  |  |  |  | M | 8 years | Epithelial | Yes |

**Supplementary Table 4:**
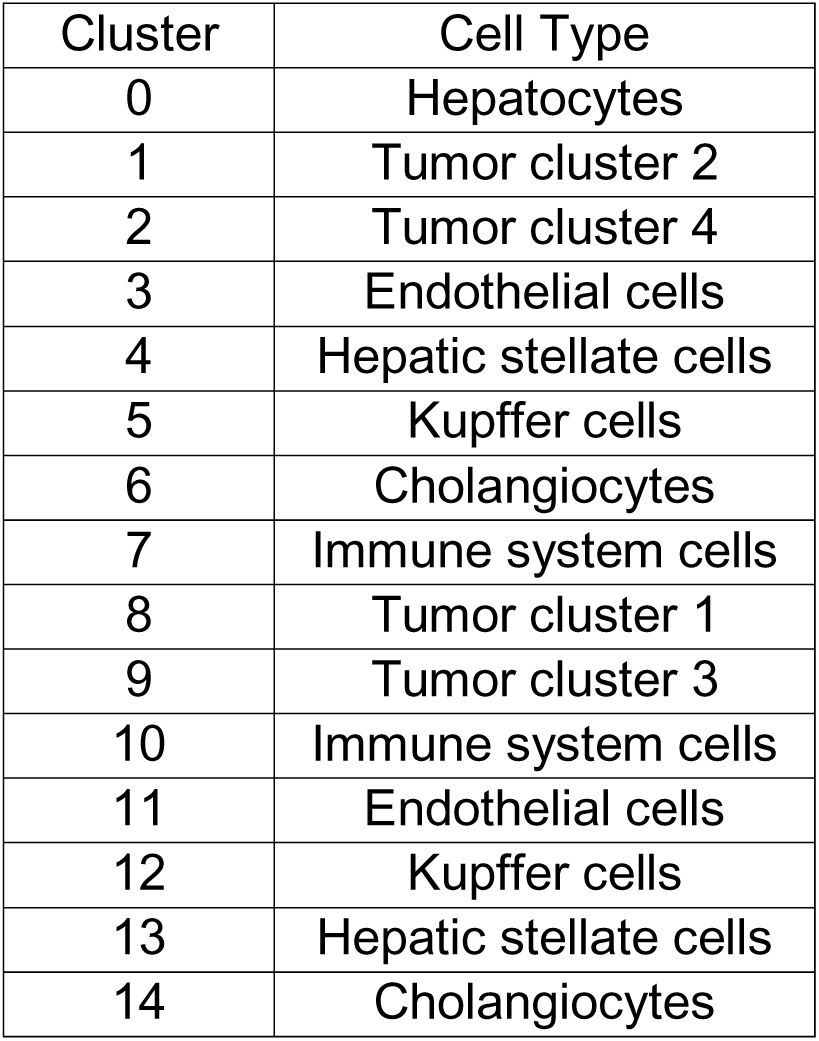
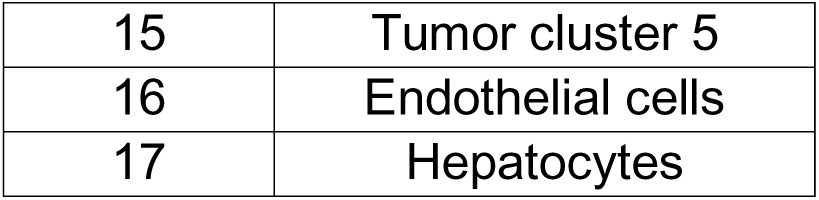
Table showing cell types represented by cluster numbers.

## Supplementary Data File Descriptions

Supplementary Data File 1 Top 50 cell markers in each cluster.csv

Supplementary Data File 2 Genelist for pathway analysis.csv

Supplementary Data File 3 Kegg and Reactome pathways enriched in cell clusters.xls

Supplementary Data File 4 Gene signature study.csv

Supplementary Data File 5 Markers_identifying_precancer_transition_cancer_states.xls

Supplementary Data File 6 List of eregulons and transcription factors (TFs) regulated in precancer, transition and cancer states of BWS and nonBWS cohorts.xlsx

